# Developmental spontaneous activity promotes sensory domains, frequency tuning and proper gain in central auditory circuits

**DOI:** 10.1101/2022.06.30.498184

**Authors:** Calvin J. Kersbergen, Travis A. Babola, Jason Rock, Dwight E. Bergles

## Abstract

Neurons that process sensory information exhibit bursts of electrical activity during development, providing early training to circuits that will later encode similar features of the external world. In the mammalian auditory system, this intrinsically generated activity emerges from the cochlea prior to hearing onset, but its role in maturation of auditory circuitry remains poorly understood. We show that selective disruption of cochlear supporting cell spontaneous activity suppressed patterned burst firing of central auditory neurons without impacting cell survival or acoustic thresholds. However, neurons within the inferior colliculus of these mice exhibited enhanced acoustic sensitivity and broader frequency tuning, resulting in wider isofrequency lamina. Despite this enhanced neural responsiveness, total tone-responsive regions of the midbrain and cortex were substantially smaller. Thus, loss of pre-hearing cochlear activity causes profound changes in neural encoding of sound, with important implications for restoration of hearing in individuals that experience disrupted activity during this critical developmental period.

## Introduction

Acoustic information is processed within discrete auditory centers of the brain to enable verbal communication, discrimination, and sound localization. The features of acoustic stimuli necessary to interpret sounds, such as frequency (pitch) and sound pressure (loudness) can be extracted from the earliest detectable sound input (Kandler, Clause and Noh, 2009; Babola *et al*., 2018), indicating that core features of auditory neural pathways are established prior to sensory experience. Before the onset of hearing, neurons within nascent sound processing networks experience periodic bouts of spontaneous electrical activity resembling future activation to pure tone acoustic stimuli (Lippe, 1994; Tritsch *et al*., 2007; Sonntag *et al*., 2009; Babola *et al*., 2018), providing the means to promote the survival of neurons in nascent auditory centers, initiate their maturation, and refine their connections through activity-dependent processes (Katz and Shatz, 1996; Kirkby *et al*., 2013; Martini *et al*., 2021). However, the role of this early patterned activity in stimulating functional maturation of auditory circuits is poorly understood, due to the inability to selectively disrupt this activity, while preserving cochlear function to allow later assessment of sound processing capabilities of neurons in the brain.

Auditory neuron burst firing is initiated peripherally in the developing cochlea when ATP is released by inner supporting cells (ISCs), which together form a transient epithelium known as Kölliker’s organ that lies adjacent to inner hair cells (IHCs). The resulting activation of purinergic P2RY1 autoreceptors on ISCs initiates a cascade of events, culminating in chloride efflux through TMEM16A (ANO-1) calcium-activated chloride channels, which draws potassium ions out of these cells to induce depolarization of nearby IHCs, triggering glutamate release and eventually burst firing of spiral ganglion neurons (SGNs) (Tritsch *et al*., 2007; Tritsch *et al*., 2010; Wang *et al*., 2015). During this pre-hearing period, projections from medial olivocochlear neurons form synapses on immature IHCs, providing efferent feedback to regulate precise burst firing patterns and bilateral representation of neural activity in the developing CNS (Clause *et al*., 2014; Wang *et al*., 2021). Although ATP release from cochlear supporting cells is stochastic and variable, it is highly effective at inducing transient, spatially restricted activation of IHCs along the length of the cochlea. As such, spontaneous events initiated in the cochlea during this period resemble future activation to pure tone acoustic stimuli, and propagate through nascent sound processing circuits to reach the auditory cortex, providing a means to reinforce connections between neurons along tonotopic boundaries throughout the auditory system (Babola *et al*., 2018). However, spontaneous activity engages many of the same components later used for sound transduction, such as glutamate release from IHCs. Manipulations that disrupt spontaneous activity by targeting these cells and components during development also result in loss of trophic support, efferent silencing, neuronal degeneration, and deafness, limiting functional *in vivo* interrogation of the auditory pathway to define the precise roles of pre-hearing spontaneous activity (Tierney, Russell and Moore, 1997; Mostafapour *et al*., 2000; McKay and Oleskevich, 2007; Noh *et al*., 2010; Clause *et al*., 2014; Tong *et al*., 2015).

To assess the role of this early patterned activity in the functional maturation of sound processing networks, we selectively disrupted the expression of TMEM16A (ANO-1) channels in ISCs and performed *in vivo* imaging of spontaneous and sound evoked neural activity in awake mice. Loss of cochlear TMEM16A channels suppressed pre-hearing neuronal calcium transients in the inferior colliculus (IC) and disturbed normally precise, tonotopically restricted activation patterns that occur at this age. Despite disruption of spontaneous activity, these animals exhibited normal cochlear structure, no cellular degeneration, or hearing loss after ear canal opening (∼postnatal day 12), allowing assessment of spatial and temporal features of sound-evoked neural activity in central auditory centers. *In vivo* imaging of neuronal calcium transients in the IC of these mice revealed that neuronal responses to pure tones were larger in amplitude over a wide range of frequencies and intensities, and more neurons were activated by a given stimulus, indicative of abnormally high gain. Moreover, neurons responded to a broader range of frequencies and the spatial map of responding neurons was wider, indicating that sharp tonotopic portioning of acoustic information was not achieved. Despite the enhanced responsiveness of auditory neurons, macroscopic imaging revealed that brain regions involved in processing sound were substantially smaller in both the midbrain and cortex. Together, these results reveal that intrinsically generated bursts of activity that emerge from the developing cochlea are required to establish appropriate sensitivity to incoming acoustic information and consolidate regions of the brain devoted to processing sound.

## Results

### TMEM16A channels are required for spontaneous activity prior to hearing onset

Opening of TMEM16A channels in ISCs results in profound chloride efflux, inducing a concomitant efflux of potassium ions that depolarizes nearby IHCs (Wang *et al*., 2015) (Fig. 1A). Our previous studies indicate that genetic inactivation of *Tmem16a* abolished most spontaneous activity in IHCs and reduced SGN burst firing. However, the *Pax2-Cre* mouse line used to inactivate *Tmem16a* exhibits incomplete recombination in the cochlea and widespread expression outside the inner ear (Wang *et al*., 2015). Moreover, it is unknown whether spontaneous burst firing is dependent on TMEM16A throughout the postnatal prehearing period (postnatal day (P) 1-P11). To determine if ISC-generated spontaneous activity requires TMEM16A throughout postnatal development, we crossed *Tmem16a^fl/fl^* mice with *Tecta-Cre* mice, which exhibit recombination within the sensory epithelium of the embryonic cochlea with limited recombination in other central auditory nuclei, enabling selective removal of TMEM16A (*Tmem16a* cKO) within supporting cells of the cochlea prior to the onset of spontaneous activity (Babola *et al*., 2020, 2021) (Fig. 1B, Supplementary Fig. 1). ISCs in *Tmem16a* cKO mice exhibited no spontaneous inward currents during whole cell patch clamp recordings throughout postnatal, prehearing development (Fig. 1C, D, Supplementary Fig. 2A-E), but retained their low membrane resistance (Supplementary Fig. 2F, G), indicative of retained gap junction coupling. Additionally, ISCs did not exhibit spontaneous osmotic crenations (Tritsch *et al*., 2007) that are induced by TMEM16A-mediated chloride efflux (Wang *et al*., 2015) (Supplementary Fig. 2H, I).

**Figure 1.**
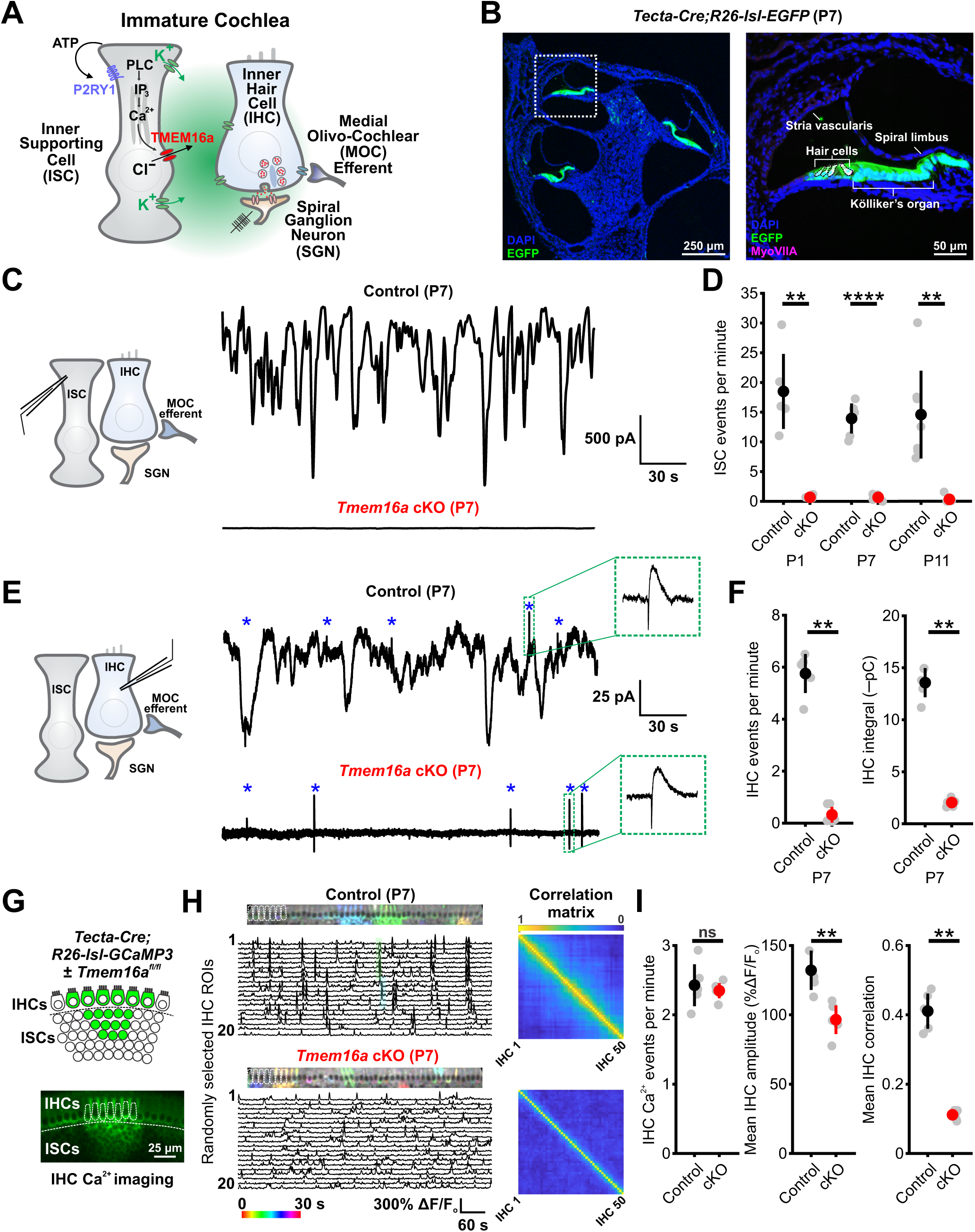
TMEM16A channels are required for spontaneous activity in the immature cochlea. (A) Mechanism of spontaneous auditory neural activity generation by inner supporting cells (ISCs) within the developing cochlea. (B) Extent of recombination of *Tecta-Cre* in cross section of the immature postnatal day 7 (P7) cochlea, as shown by EGFP (green) reporter expression. White square indicates site of high magnification. Inner and outer hair cells are labeled by immunoreactivity to Myosin VIIa (MyoVIIa, magenta; circled by black dashed lines). See Supplementary Fig. 1 for additional characterization of *Tecta-Cre*. (C) Whole cell patch clamp recordings of spontaneous activity from P7 control (*Tmem16a^fl/fl^*) and P7 *Tmem16a* cKO (*Tecta-Cre;Tmem16a^fl/fl^*) ISCs within acutely isolated cochleae. (D) Quantification of spontaneous inward current frequency in ISCs at three ages (P1, P7, P11) encompassing postnatal development. Gray circles represent individual animals, colored circles represent the mean value and error bars represent standard deviation. n = 5, 6, 7 control ISCs, n = 6, 6, 7 *Tmem16a* cKO ISCs (P1, P7, P11); p = 0.0043, 4.1330e-7, 0.0061 (P1, P7, P11), Wilcoxon rank sum test (P1, P11) or two-sample t-test (P7) with Benjamini-Hochberg correction for multiple comparisons. (E) Whole cell patch clamp recordings of spontaneous activity from P7 control and P7 *Tmem16a* cKO inner hair cells (IHCs) within acutely isolated cochleae. Asterisks indicate medial olivocochlear (MOC) efferent synaptic response (inset) present in both control and *Tmem16a* cKO recordings. (F) Quantification of spontaneous inward current frequency and integral (charge transfer) in IHCs at P7. n = 5 control IHCs, 6 *Tmem16a* cKO IHCs; p = 0.0043, 0.0043 (frequency, integral), Wilcoxon rank sum test with Benjamini-Hochberg correction. (G) Schematic depicting calcium imaging in IHCs within excised cochlea that express a Cre-dependent genetically encoded calcium indicator GCaMP3. ROIs are placed over each IHC. (H) (left) Pseudocolored projection of IHC spontaneous calcium activity over 30 s and corresponding raster plot over 10 minutes in control (*Tecta-Cre;Tmem16a^fl/+^;R26-lsl-GCaMP3*) and *Tmem16a* cKO (*Tecta-Cre;Tmem16a^fl/fl^;R26-lsl-GCaMP3*) cochleae. (right) Mean correlation matrix of calcium activity between individual IHCs. n = 5 control, 6 *Tmem16a* cKO cochleae. (I) Quantification of spontaneous calcium transients in IHCs. n = 5 control, 6 *Tmem16a* cKO cochleae; p = 0.7922, 0.0065, 0.0065 (frequency, amplitude, correlation), Wilcoxon rank sum test with Benjamini-Hochberg correction.

Consistent with the critical role of TMEM16A in inducing local extracellular potassium transients, IHCs in P7 cochleae of *Tmem16a* cKO mice no longer exhibited the slow inward currents that induce bursts of calcium action potentials (Beutner and Moser, 2001; Tritsch *et al*., 2010; Johnson *et al*., 2011) (Fig. 1E, F), leaving only infrequent efferent synaptic responses (Glowatzki and Fuchs, 2000) (Fig. 1E, *asterisks* and *inset*). Unexpectedly, calcium imaging in *Tmem16a* cKO mice revealed the presence of spontaneous calcium transients in IHCs (Fig. 1G, H); however, these transients were no longer coordinated among neighboring IHCs and exhibited reduced amplitudes (Fig. 1H, I). Moreover, calcium transients in *Tmem16a* cKO ISCs no longer correlated with local IHC calcium activity (Supplementary Fig. 3E-H). These isolated IHC calcium events arise from tonic IHC depolarization, due to accumulation of extracellular potassium around IHCs when TMEM16A-mediated osmotic crenation is prevented (Babola *et al*., 2020) (Supplementary Fig. 4A-D). Notably, this phenotype recapitulates IHC behavior in mice in which ATP-dependent activation of ISCs was prevented (Babola *et al*., 2020, 2021). Similarly, SGNs in *Tmem16a* KO mice exhibit altered firing patterns, including a near elimination of long burst events and prolongation of inter-spike intervals (Wang *et al*., 2015). A recent report suggested that TMEM16A enhances ATP release through positive feedback mechanisms in the cochlea (Maul *et al*., 2022); however, we observed no significant change in the frequency, amplitude, duration, or area of calcium transients in ISCs of *Tmem16a* cKO mice (Supplementary Fig. 3A-D, Supplementary Movie 1), in accordance with prior observations that TMEM16A removal does not disrupt ATP release or the ability of purinergic receptors to mobilize intracellular calcium stores (Fig. 1A) (Wang *et al*., 2015). Together, these results indicate that TMEM16A is required to induce periodic, coordinated activation of IHCs throughout the postnatal pre-hearing period, providing a means to selectively disrupt spontaneous activity in the auditory system during this crucial period of development.

### Deletion of *Tmem16a* suppresses burst firing of central auditory neurons prior to hearing onset

ISC-induced depolarization of IHCs triggers excitation of SGNs, leading to discrete bursts of action potentials that propagate throughout the central auditory system (Sonntag *et al*., 2009; Tritsch *et al*., 2010; Clause *et al*., 2014). Each ATP release event results in coordinated IHC activation within a restricted region of the cochlea, enabling correlated firing of neurons within isofrequency lamina that will ultimately process similar frequencies of sound. To determine how disruption of TMEM16A-induced ionic flux in the cochlea influences spontaneous activity patterns of central auditory neurons *in vivo*, we performed macroscopic *in vivo* imaging of the IC in awake, unanesthetized mice expressing the genetically encoded Ca^2+^ indicator GCaMP6s in neurons (*Snap25-T2A-GCaMP6s*) (Madisen *et al*., 2015) (Fig. 2A). In control mice (*Tmem16a^fl/fl^*;*Snap25-T2A-GCaMP6s*) (P7), neurons within isofrequency lamina exhibited periodic calcium increases, resulting in spatially restricted bands of activity that were mirrored across both lobes of the IC (Babola *et al*., 2018; Wang *et al*., 2021) (Fig. 2B; *Events 1-4*). The spontaneous activity of auditory neurons was dramatically suppressed in *Tmem16a* cKO mice, with fewer calcium transients detected in the IC (Fig. 2C, D, Supplementary Movie 2). Residual activity in these mice consisted of low amplitude, brief events (Fig. 2C; *Events 1, 2*), interspersed with large amplitude, long-duration events occurring once every ∼2 minutes that were no longer confined to narrow isofrequency boundaries and displayed high correlation between IC lobes (Fig. 2C, D; *Events 3, 4*). These changes in activity are reflected in the histogram of event amplitudes (Fig. 2E) and cumulative distribution of durations (Fig. 2F), with *Tmem16a* cKO mice exhibiting shifts towards the distribution extremes for both parameters. Analysis of fluorescence changes across the tonotopic axis revealed that spontaneous events in *Tmem16a* cKO mice resulted in a broader activation of neurons (Fig. 2G-I), with a significant increase in the mean spatial activation across animals (control, 250.0 ± 25.4 µm; cKO, 325.6± 41.4 µm; p = 0.002, two-sample t-test, n = 6 and 8 mice, respectively), indicating that loss of cochlear TMEM16A expression disrupts the precise patterning of correlated neuronal calcium activity within isofrequency lamina. Importantly, retina-induced spontaneous waves of neuronal activity in the superior colliculus (Ackman, Burbridge and Crair, 2012) did not exhibit changes in frequency or duration in *Tmem16a* cKO mice (Supplementary Fig. 5), suggesting that cochlear deletion of *Tmem16a* did not induce global physiological changes within the CNS.

**Figure 2.**
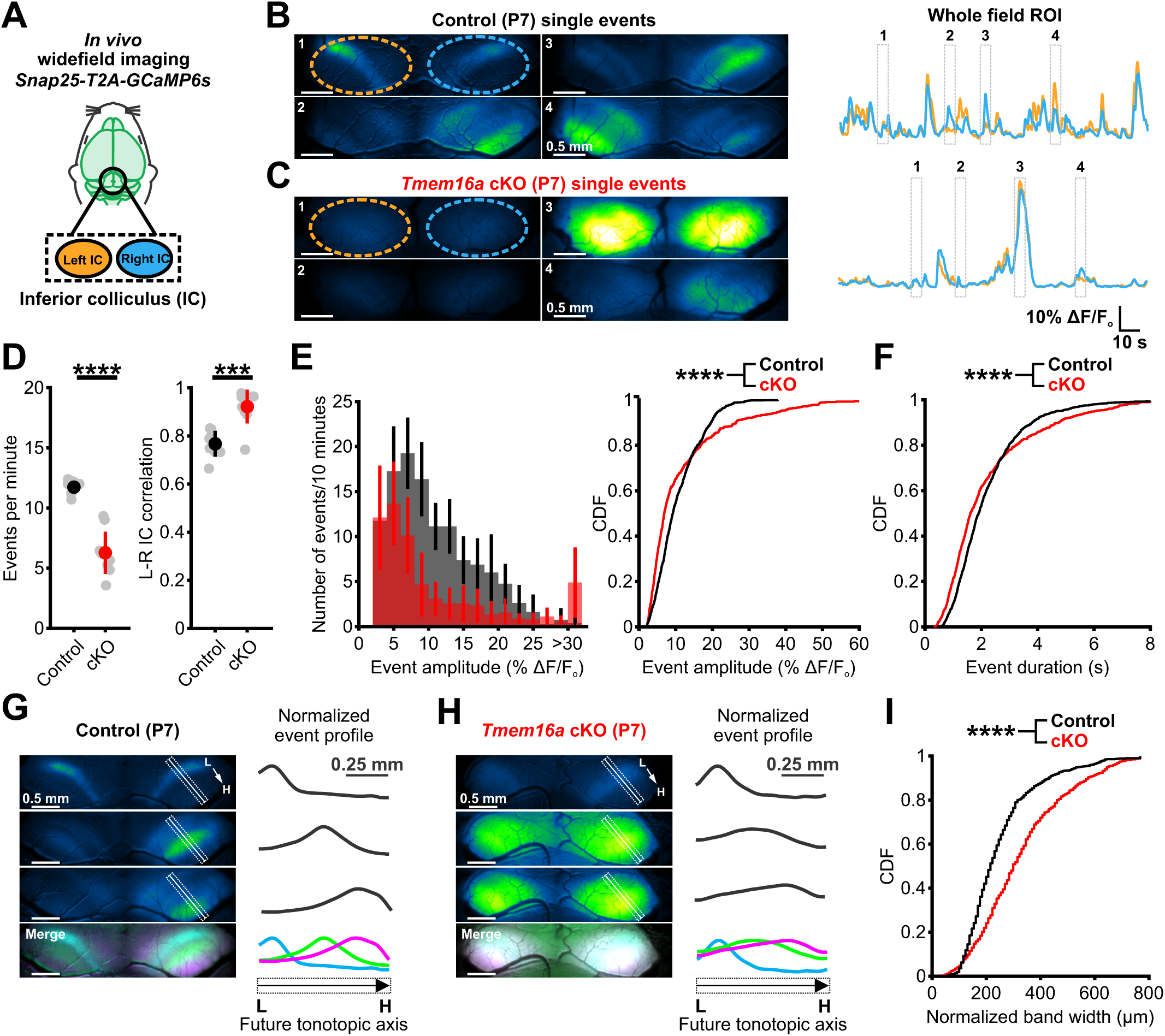
Deletion of *Tmem16a* suppresses calcium transients in central auditory neurons prior to hearing onset. (A) *In vivo* widefield imaging paradigm to visualize spontaneous neural activity in the inferior colliculus (IC) of unanesthetized mouse pups. (B) (left) Four examples of calcium transients in the IC from P7 control (*Tmem16a^fl/fl^;Snap25-T2A-GCaMP6s*) mouse. Colored dashed circles denote left and right ROIs. (right) Fluorescence trace over time of spontaneous activity in left (orange) and right (blue) IC with example single events highlighted. (C) Same as (B), but from P7 *Tmem16a* cKO (*Tecta-Cre;Tmem16a^fl/fl^;Snap25-T2A-GCaMP6s*) mouse. (D) Quantification of spontaneous event frequency and left-right IC correlation. n = 8 control mice, 9 *Tmem16a* cKO mice; p = 1.0033e-5, 2.6549e-4 (frequency, correlation), two-sample t-test with unequal variances (frequency) or Wilcoxon rank sum test (correlation) and Benjamini-Hochberg correction. (E) (left) Histogram of spontaneous event amplitude. (right) Cumulative distribution of spontaneous event amplitude. n = 940 events from 8 control mice, 560 events from 9 *Tmem16a* cKO mice; p = 6.4020e-11, two-sample Kolmogorov-Smirnov test with Benjamini-Hochberg correction. (F) Cumulative distribution of spontaneous event duration. n = 940 events from 8 control mice, 560 events from 9 *Tmem16a* cKO mice; p = 6.373e-6, two-sample Kolmogorov-Smirnov test with Benjamini-Hochberg correction. (G) (left) Calcium transients in control mice within restricted bands along the future tonotopic axis (low (L) to high (H) frequency) of the IC in P7 control mouse. White box indicates averaged region (along short axis) to generate spatial tonotopic fluorescence profile. Pseudocolored merged image highlights distinct activation domains by each event. (right) Plot of spatial fluorescence profile along future tonotopic axis, normalized to peak fluorescence change, for the spontaneous event to left. (H) Same as (G), but from a P7 *Tmem16a* cKO mouse. (I) Cumulative distribution of normalized band width (75^th^ percentile) for spontaneous events. n= 986 events from 6 control mice, 550 events from 8 *Tmem16a* cKO mice; p = 1.6325e-23, two-sample Kolmogorov-Smirnov test.

*In vivo* two-photon imaging of spontaneous neural activity in the IC at this age confirmed that activity patterns observed during widefield imaging arise from local neuronal activity (including projections) and not light scattering from other brain regions. Automated grid-based analyses revealed that IC neurons in *Tmem16a* cKO mice exhibited significantly fewer correlated events, but when they occurred, they typically encompassed a much larger portion of the IC (Supplementary Fig. 6). To determine if these changes in IC spontaneous activity patterns occur throughout the pre-hearing period, we performed widefield imaging from mice aged P10-P11, just prior to ear canal opening (Mikaelian and Ruben, 1965; Geal-Dor *et al*., 1993; Anthwal and Thompson, 2016). *Tmem16a* cKO mice at this age continued to exhibit reduced activity and dramatic increases in the amplitude and spatial width of residual spontaneous events (Supplementary Fig. 7), consistent with previous studies indicating that the cochlea uses a similar mechanism to induce spontaneous activity throughout early development (Tritsch and Bergles, 2010; Babola *et al*., 2021). Together, these results show that disruption of ion flux from ISCs alters the normal spatial and temporal patterning of spontaneous neural activity in the central auditory system prior to hearing onset.

### Cochlear structure and acoustic sensitivity are preserved in *Tmem16a* cKO mice

To determine how pre-hearing spontaneous activity influences the functional organization of central auditory centers, the ability of the cochlea to transduce acoustic stimuli must be intact after knockout of *Tmem16a*. TMEM16A expression rapidly declines after ear canal opening (Supplementary Fig. 8A), suggesting that it should primarily contribute to intrinsically-generated, rather than sound-evoked activity. To determine if developmental deletion of *Tmem16a* alters cochlear structure or cell survival, we performed histological assessments of *Tmem16a* cKO cochleae at P14, just after hearing onset, and at P21, when mice exhibit normal hearing thresholds. No changes in gross structure of the cochlea, organ of Corti, or tectorial membrane were observed in these mice (Fig. 3A), and no degeneration of hair cells or SGNs was evident across the length of the cochlea at either P14 or P21 (Fig. 3B-E, Supplementary Fig. 8B-F). To determine if the ability to transduce acoustic stimuli is intact in mice with disrupted spontaneous activity, we measured auditory brainstem responses (ABRs) at P14 and P21. Auditory thresholds to click and pure tone stimuli were unaltered in *Tmem16a* cKO mice (Fig. 3F, G, Supplementary Fig. 8G-I), although subtle differences in ABR waveforms were evident, including a slightly lower amplitude (Fig. 3H) and longer latency of wave 1 and subsequent waveforms (Fig. 3I), suggesting that SGN activation may be slightly less synchronous. Together, these results indicate that the cochlea retains the ability to transduce sound and convey neural impulses to the CNS despite the loss of TMEM16A and disruption of spontaneous activity prior to hearing onset.

**Figure 3.**
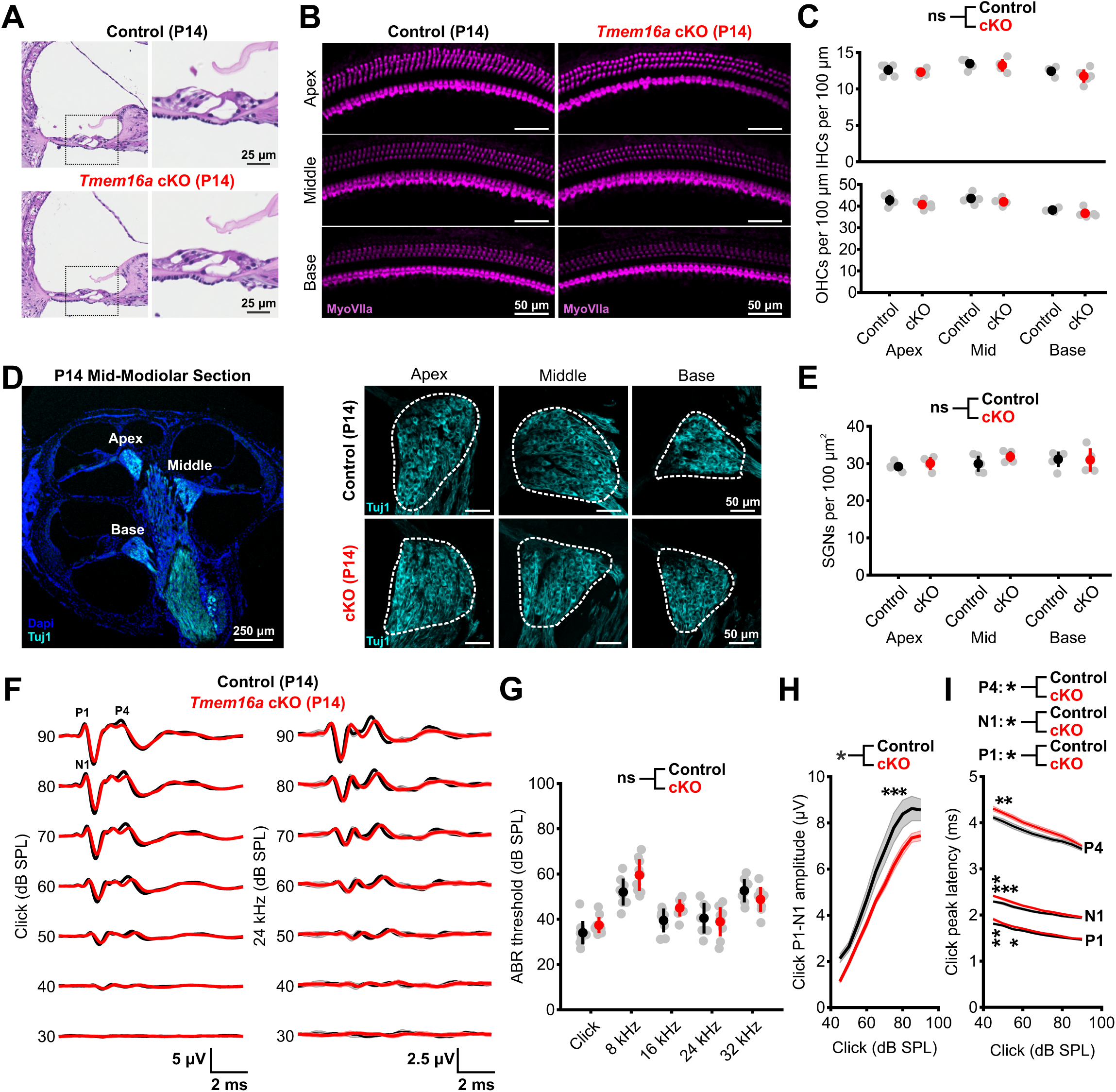
Cochlear structure and sound detection are preserved in *Tmem16a* cKO mice. (A) Hematoxylin and eosin stain of cross sections from P14 control (*Tmem16a^fl/fl^*) and *Tmem16a* cKO (*Tecta-Cre;Tmem16a^fl/fl^*) basal cochleae. Black square indicates region of high magnification. (B) Hair cells in whole mount preparations of apical, middle, and basal P14 control and *Tmem16a* cKO cochleae labeled by immunoreactivity to MyoVIIa (magenta). (C) Quantification of inner hair cell (IHC) and outer hair cell (OHC) density. n = 6 cochleae from 6 control mice, 5 cochleae from 5 *Tmem16a* cKO mice; p = 0.2392, 0.0840 (IHCs, OHCs), linear mixed model. (D) (left) Low magnification mid-modiolar cross section of P14 cochlea with spiral ganglion neurons (SGNs) labeled by immunoreactivity to β-III tubulin (Tuj1, cyan). (right) High magnification images of SGN soma in apical, middle, and basal regions of the cochlea in P14 control and *Tmem16a* cKO mice. (E) Quantification of SGN density. n = 5 cochleae from 5 control mice, 4 cochleae from 4 *Tmem16a* cKO mice; p = 0.5057, linear mixed model. (F) Mean auditory brainstem response (ABR) trace to click (left) or 24 kHz pure tone pip (right) stimuli from 90 to 20 dB sound pressure level (SPL) in P14 control and *Tmem16a* cKO mice. P1= ABR peak 1, N1 = ABR trough 1, P4 = ABR peak 4. (G) Quantification of ABR threshold to click and pure tone stimuli. n = 9 control mice, 10 *Tmem16a* cKO mice; p = 0.0962, linear mixed model. (H) Quantification of click ABR wave 1 (P1:N1) amplitude across a range of sound levels. n = 9 control mice, 10 *Tmem16a* cKO mice; mean ± SEM, p = 0.0143, linear mixed model with Sidák post hoc test. (I) Quantification of click ABR wave latency (P1, N1, P4) across a range of sound levels. n = 9 control mice, 10 *Tmem16a* cKO mice; mean ± SEM, * p = 0.0203, 0.0358, 0.0305 (P1, N1, P4), linear mixed model with Sidák post hoc test.

### Suppression of pre-hearing spontaneous activity enhances the gain of central auditory neurons in awake mice

To determine how auditory neurons in the CNS respond to sound when deprived of normal burst firing prior to hearing onset, we assessed sound-evoked neural activity within the IC just after ear canal opening (P13-P15), using widefield imaging of GCaMP6s in awake, unanesthetized mice (Fig. 4A). In both control and *Tmem16a* cKO mice, presentation of pure tones to the left ear elicited broad regions of neural activity in both lobes of the IC that were aligned to diagonally-oriented isofrequency domains, with the strongest response observed in the contralateral IC (Babola *et al*., 2018) (Fig. 4B). Low frequency tones elicited a single band of activity in the central portion of the IC, while higher frequency tones elicited dual bands in more lateral regions of the IC (Fig. 4B), following the known spatial segregation of frequency information within the IC (Stiebler and Ehret, 1985; Barnstedt *et al*., 2015; Wong and Borst, 2019). Moreover, both control and *Tmem16a* cKO mice exhibited similar response thresholds (Fig. 4C, D), in accordance with ABR measurements (Fig. 3G), indicating that core aspects of auditory encoding remain intact. However, *Tmem16a* cKO mice exhibited several anomalies, including a marked increase in the amplitude of sound-evoked responses to a given sound intensity, particularly to suprathreshold tones above 3 kHz (Fig. 4E, F). This increase in gain was accompanied by a widening of the area responsive to individual tones (e.g. band width) along the dorsal tonotopic axis across a range of frequencies and sound intensities (Fig. 4G, H, Supplementary Fig. 9A-J), resulting in increased overlap of pure tone isofrequency lamina and indicating that suppression of auditory spontaneous activity during the pre-hearing period results in inappropriately high gain within sound processing circuits.

**Figure 4.**
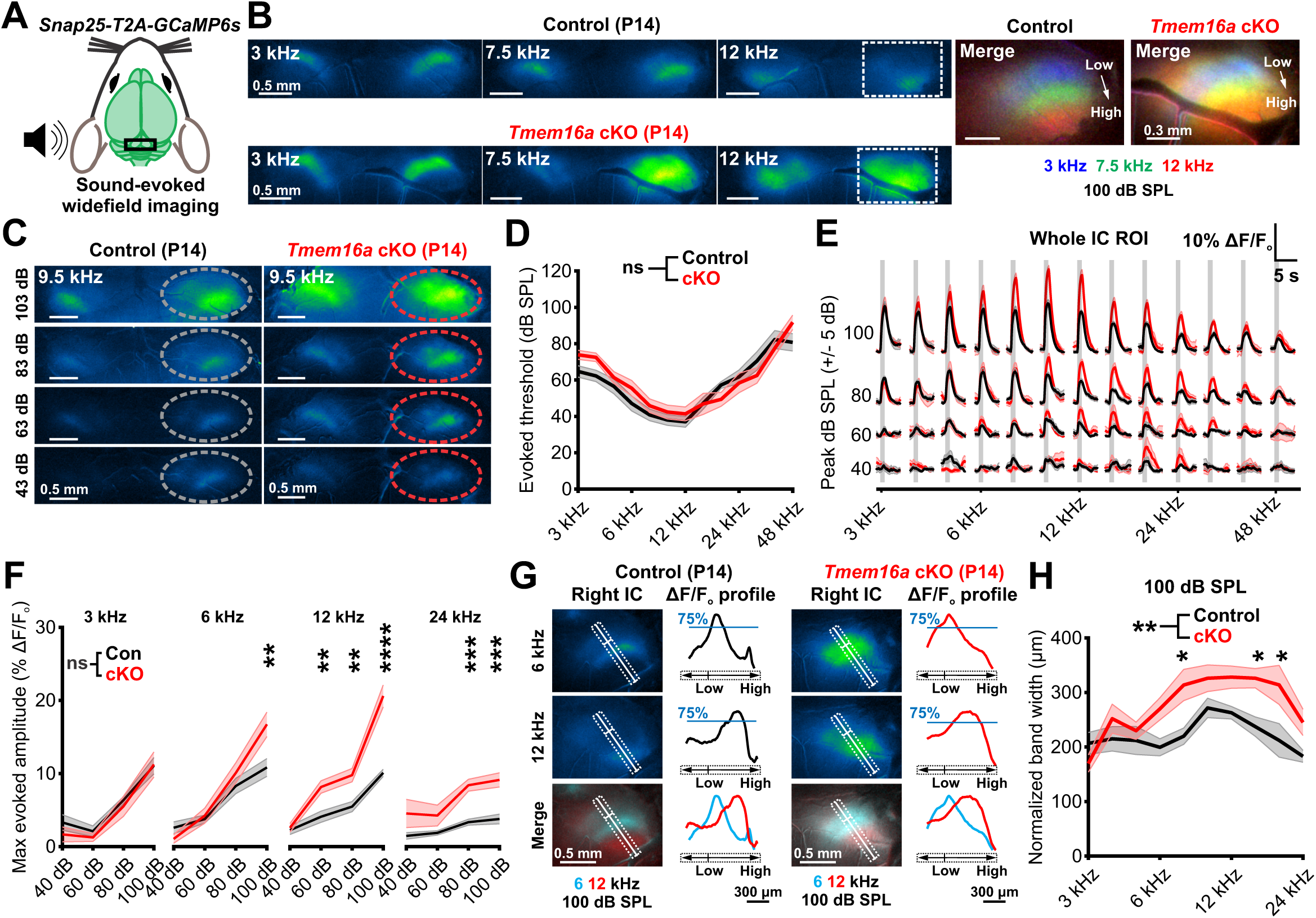
Suppression of pre-hearing spontaneous activity enhances the gain of central auditory neurons. (A) *In vivo* widefield imaging of tone-evoked inferior colliculus (IC) neural activity in unanesthetized mice after hearing onset. (B) Tone-evoked neural calcium transients in IC from P14 control (*Tmem16a^fl/fl^;Snap25-T2A-GCaMP6s*) and P14 *Tmem16a* cKO (*Tecta-Cre;Tmem16a^fl/fl^;Snap25-T2A-GCaMP6s*) mice at 100 dB SPL. Merged and zoomed image of right IC (right) shows tonotopic segregation of pseudocolored pure tone responses; cyan and yellow colors indicate areas of overlap. (C) IC neural calcium transients to a 9.5 kHz stimulus from 103 to 43 dB sound pressure level (SPL) in a control and *Tmem16a* cKO mouse. Circle in right IC depicts ROI for subsequent quantification of threshold and amplitude. (D) Quantification of pure tone sound-evoked thresholds. n = 11 control mice, 8 *Tmem16a* cKO mice; p = 0.0941, repeated measures ANOVA with lower bound p value adjustment. (E) Quantification of tone-evoked fluorescence in IC across a range of frequency and sound level stimuli in control and *Tmem16a* cKO mice. Vertical gray bar indicates tone presentation. N = 9-12 control mice, 5-10 *Tmem16a* cKO mice, mean ± SEM. (F) Rate-level functions characterizing maximum whole IC response amplitude at 3, 6, 12, and 24 kHz. n = 9-12 control mice, 5-10 *Tmem16a* cKO mice; linear mixed model with Sidák post hoc test. (G) Measurement of pure tone evoked spatial activation (75^th^ percentile) normalized to peak fluorescence response amplitude along the tonotopic axis of the IC at 100 dB SPL. (H) Quantification of spatial evoked fluorescence along tonotopic axis at 100 dB SPL. n = 9 control mice, 8 *Tmem16a* cKO mice; mean ± SEM, p = 0.0057, linear mixed model with Sidák post hoc test.

### Neurons display increased gain and broader frequency sensitivity when deprived of spontaneous burst firing

The increased acoustic sensitivity visible through widefield imaging in *Tmem16a* cKO mice could arise from larger calcium increases within individual cells or from increased numbers of neurons responding to pure tones. To assess the response properties of individual auditory neurons, we performed two photon calcium imaging within the central nucleus of the IC (Fig. 5A). Both neuronal somata and neuropil responsive to a given frequency exhibited markedly larger increases in calcium to tone presentation in *Tmem16a* cKO mice (Fig. 5B, C, Supplementary Fig. 10A-F, Supplementary Movie 3), indicating that these gain changes manifest within single neurons. Additionally, more neurons within each field responded to each tone (Fig. 5D), demonstrating that these changes in sensitivity were pervasive within sound-responsive collicular neurons. We also determined the frequency tuning of IC neurons by measuring their response to a range of frequencies (3-24 kHz) at different sound attenuation levels (40-90 dB SPL). Neurons in control mice exhibited sharp tuning, responding to a narrow range of frequencies, with calcium levels increasing with higher sound intensities (Fig. 5E, F). In contrast, neurons in *Tmem16a* cKO mice exhibited much broader tuning, responding to a wider range of frequencies (Fig. 5E-H, Supplementary Fig. 10G-I, Supplementary Movie 4), with a significant increase in the mean maximum bandwidth across animals (control, 0.76 ± 0.17 octaves; cKO, 1.48 ± 0.31 octaves; p =7.7804e-5, two-sample t-test, n = 8 and 7 mice), and exhibited larger calcium increases to their best frequency stimulus at a given sound intensity (Fig. 5I, Supplementary Fig. 10J). Thus, individual auditory neurons exhibit both higher gain and broader tuning when deprived of spontaneous burst firing prior to hearing onset.

**Figure 5.**
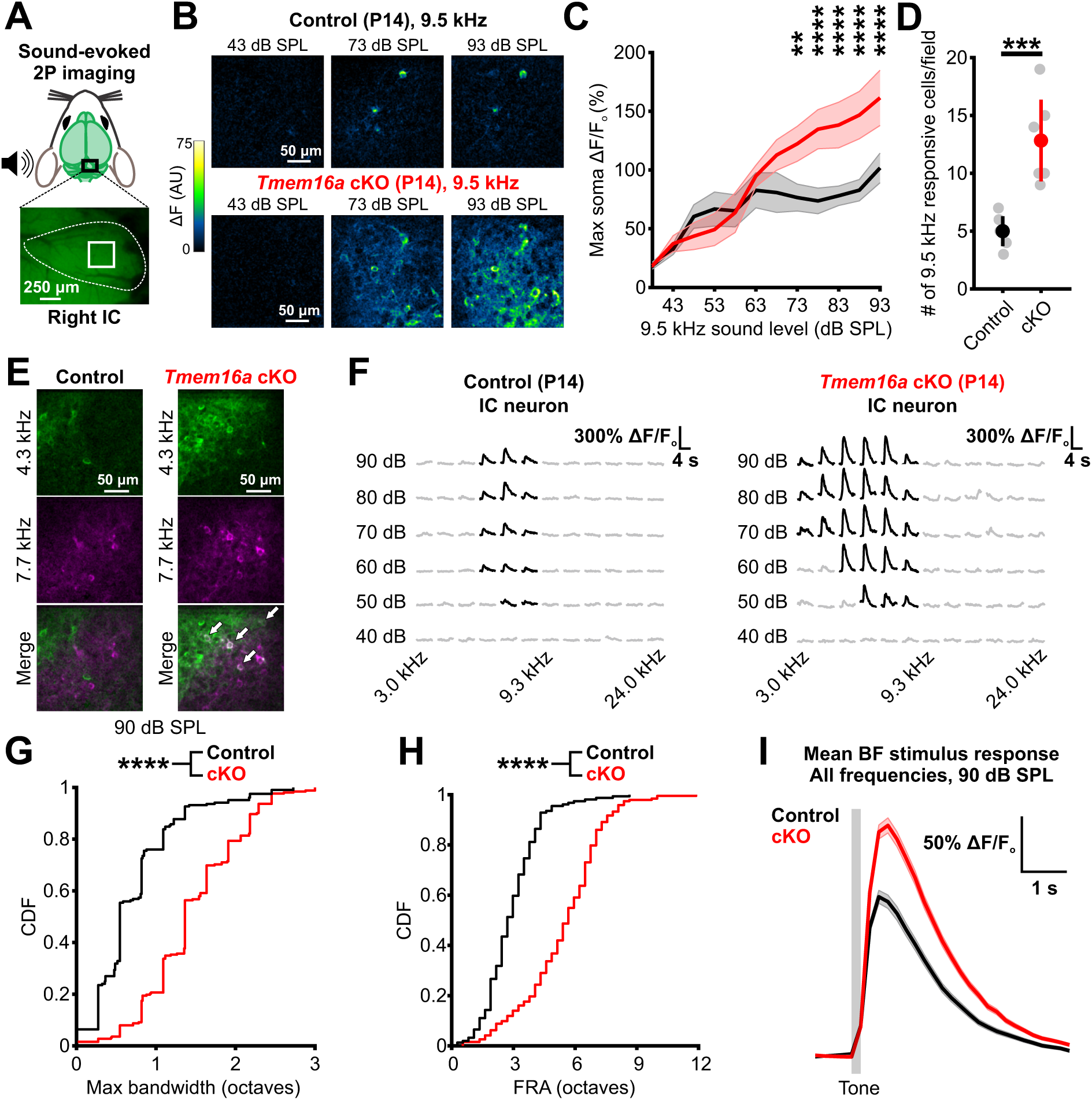
Disrupted patterns of pre-hearing spontaneous activity elicit broadening of receptive fields. (A) Schematic depicting site of two photon (2P) imaging in the central nucleus of the inferior colliculus in P14 mice. (B) Two-photon high magnification imaging of tone-evoked fluorescence to increasing sound levels in IC neurons and neuropil in P14 control (*Tmem16a^fl/fl^;Snap25-T2A-GCaMP6s*) and *Tmem16a* cKO (*Tecta-Cre;Tmem16a^fl/fl^;Snap25-T2A-GCaMP6s*) mice. (C) Fluorescence changes within responsive neuronal soma elicited by increasing intensity of a 9.5 kHz pure tone. n = 6 control mice (30 cells), 6 *Tmem16a* cKO mice (77 cells); mean ± SEM, p = 0.0417, linear mixed model with Sidák post hoc test. (D) Quantification of the number of responsive neurons to a 9.5 kHz pure tone within the fixed imaging field. n = 6 control mice, 6 *Tmem16a* cKO mice; p = 8.9678e-4, two-sample t-test with unequal variances. (E) Pseudocolored tone-evoked fluorescence in IC neurons at 90 dB SPL in a control and *Tmem16a* cKO mouse. White arrows indicate neurons responsive to both stimuli in merged image. (F) Fluorescence changes in a representative IC neuron to a range of frequency and intensity stimuli in a control and *Tmem16a* cKO mouse. Black traces represent positive responses to the stimulus, while gray traces indicate unresponsiveness to the stimulus. (G) Quantification of maximum frequency bandwidth of IC neurons at any sound level. n = 207 control cells (8 mice), 252 *Tmem16a* cKO cells (7 mice); p = 4.7586e-32, two-sample Kolmogorov-Smirnov test. (H) Cumulative distribution of frequency response area (FRA) of tone-responsive neural soma in the IC. n = 157 control cells (6 mice), 193 *Tmem16a* cKO cells (4 mice); p = 1.2817e-31, two-sample Kolmogorov-Smirnov test. (I) Mean fluorescence change in response to a neuron’s best frequency stimulus at 90 dB SPL. Vertical bar indicates tone presentation. n = 235 control cells, 295 *Tmem16a* cKO cells; mean ± SEM.

### Spatial compression of frequency representation after disruption of developmental spontaneous activity

To determine if spontaneous activity influences the partitioning of frequency information within the IC (i.e. the tonotopic map), we also measured the spatial segregation of tone-evoked responses across the IC, perpendicular to the isofrequency lamina (Fig. 6A). The peak response location to supratheshold (100 dB SPL), pure tone stimuli (i.e. site of best frequency) shifted from the center for low frequencies, to the lateral regions of the IC for higher frequencies in both control and *Tmem16a* cKO mice (Fig. 6A, B, *red and black dashed lines*), indicating that gross tonotopic partitioning is maintained. However, the location of the peak response shifted significantly less with increasing frequency in *Tmem16a* cKO mice (Fig. 6B, C); in concert with the broadening of frequency responsiveness, there was less spatial separation between frequency responsive domains and overlapping pure tone response regions (Fig. 6A, D, Supplementary Movie 5). This macroscopic spatial compression of the tonotopic map persisted over a wide range of sound intensities (60 and 80 dB SPL) (Supplementary Fig. 11A-G), suggesting that a fundamental change in frequency encoding occurs upon disruption of prehearing spontaneous activity. Low magnification, two photon imaging of neurons in the central nucleus of the IC provided further evidence of this dramatic phenomenon, as a much broader distribution of frequencies at suprathreshold levels were represented within the fixed area of the imaging field (Fig. 6E). Furthermore, maps of neuronal best frequency (frequency that elicits the largest calcium response) and center frequency (frequency of calcium response at lowest sound intensity) obtained with higher magnification imaging revealed that individual regions of the IC contained neurons with a much wider range of best and center frequencies (Fig. 6F-I), confirming an underlying topographic organizational change beyond expansion of neuronal receptive fields. Together, these data indicate that the area devoted to processing sound information within the IC is compressed when input from the cochlea is suppressed during the postnatal, prehearing period.

**Figure 6.**
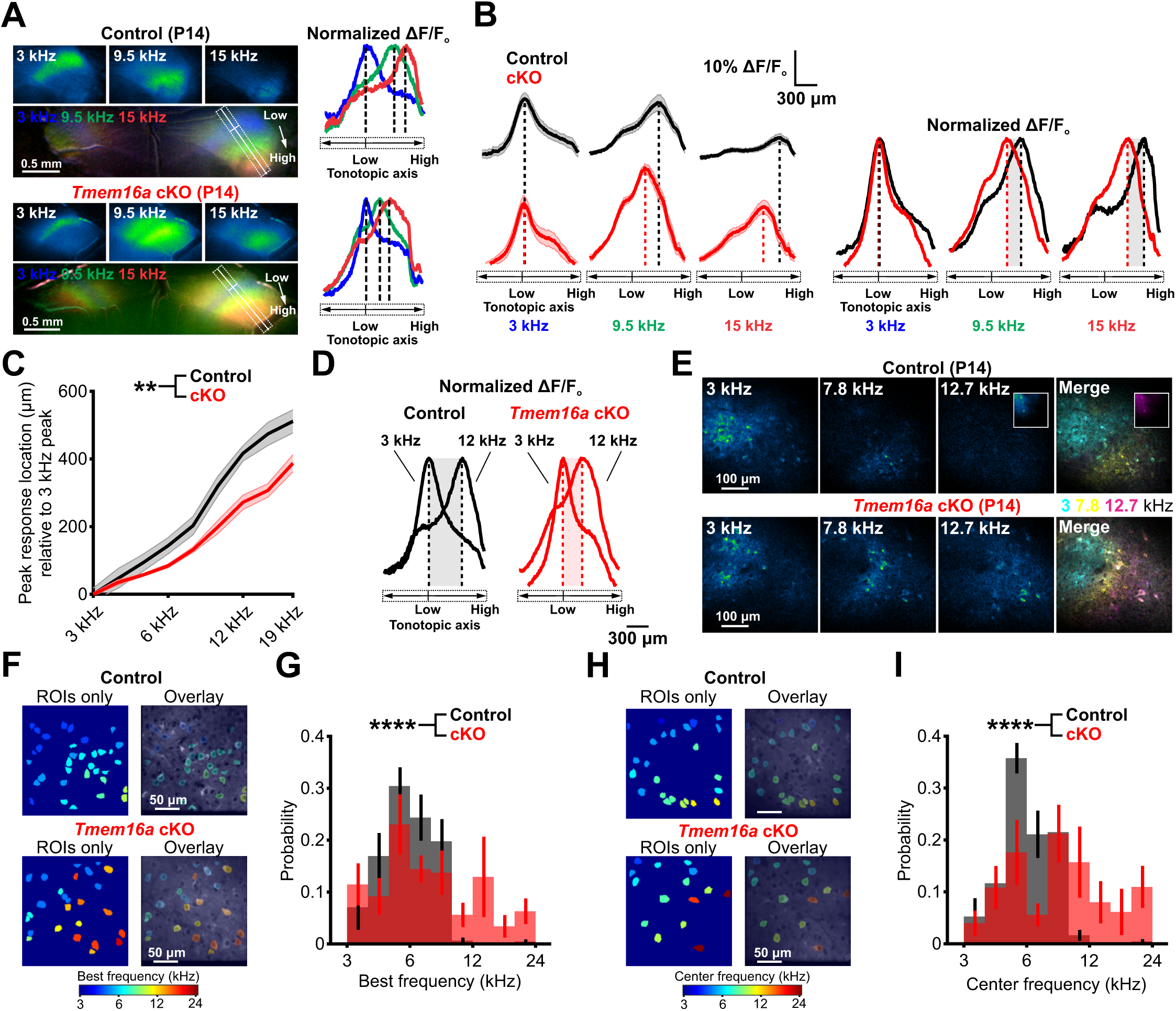
Spatial compression of frequency representation after disruption of developmental spontaneous activity. (A) (left) Pseudocolored individual and merged images of tone-evoked neural calcium transients in P14 control (*Tmem16a^fl/fl^;Snap25-T2A-GCaMP6s*) and P14 *Tmem16a* cKO (*Tecta-Cre;Tmem16a^fl/fl^;Snap25-T2A-GCaMP6s*) mice. Rectangular ROIs were placed along the tonotopic axis of the contralateral IC (low to high frequency), perpendicular to pure tone evoked bands, to determine the peak response location. (right) Plot of normalized pure tone fluorescence response along tonotopic axis for animal at left, with dashed lines indicating peak response location. (B) (left) Plot of mean tone-evoked fluorescence along the tonotopic axis of the IC in control (top) and *Tmem16a* cKO (bottom). Dashed line indicates location of peak response along tonotopic axis. (right) Normalized mean fluorescence along the tonotopic axis of the IC. Gray shading indicates shift in peak response location between control and *Tmem16a* cKO mice. n = 9 control mice, 8 *Tmem16a* cKO mice, mean ± SEM. (C) Quantification of peak response location relative to 3 kHz (lowest frequency) peak response location. n = 9 control mice, 8 *Tmem16a* cKO mice; mean ± SEM, p = 0.00540, linear mixed model. (D) Plot of mean normalized tone-evoked fluorescence along the tonotopic axis of the IC to 3 kHz and 12 kHz pure tones. Dashed line indicates peak response location, shading indicates difference between 3 kHz and 12 kHz peak location. n = 9 control mice, 8 *Tmem16a* cKO mice. (E) Low magnification two-photon imaging of tone-evoked neural activity in the IC of a P14 control and *Tmem16a* cKO mouse at a z-depth of 150 μm. No 12.7 kHz responsive cells were observed within the control field of view, but were detected 150 μm laterally and 50 μm deeper than the field of view (inset). Pseudocolored merged image depicts spatial separation of tonal responses of neurons and neuropil. (F) High magnification two-photon imaging of IC neuronal ROIs, pseudocolored by best frequency (BF), in a P14 control and *Tmem16a* cKO mouse. (G) Histogram of BF distribution within a high magnification IC field of view. n = 207 control cells (8 mice), 252 *Tmem16a* cKO cells (7 mice); p = 7.1869e-5, two-sample Kolmogorov-Smirnov test. (H) High magnification two-photon imaging of IC neuronal ROIs, pseudocolored by center frequency (CF), in a P14 control and *Tmem16a* cKO mouse. (I) Histogram of CF distribution within a high magnification IC field of view. n = 207 control cells (8 mice), 252 *Tmem16a* cKO cells (7 mice); p = 1.1214e-10, two-sample Kolmogorov-Smirnov test.

Neuronal activity that is initiated in the cochlea prior to hearing onset propagates throughout the central auditory system, reliably inducing correlated firing of neurons within isofrequency zones in the primary auditory cortex (AC). However, dual IC-AC imaging revealed that approximately half of burst events in AC arise from non-cochlear sources, raising questions about the distinct roles of peripheral and central activity sources in shaping auditory cortical development (Siegel *et al*., 2012; Babola *et al*., 2018; Gribizis *et al*., 2019). To determine how disruption of spontaneous peripheral input influences later sound processing within AC, we performed widefield imaging of sound evoked responses in P13-P16 mice. Auditory cortex is segregated into discrete high and low frequency responsive regions, or foci, along multiple tonotopic gradients (Fig. 7A, *red and blue areas*) (Issa *et al*., 2014; Romero *et al*., 2020). Remarkably, in contrast to the enhanced gain observed in IC, the amplitudes of neuronal responses in A1 were comparable in control and *Tmem16a* cKO mice after hearing onset (P13-P14) (Fig. 7B, C). However, the spatial separation between frequencies (6-24 kHz, 80 dB SPL) was reduced (Fig. 7D, E), with a change in the spatial representation of higher frequency tones; rather than split into two discrete foci as seen in control mice, 24 kHz responses in *Tmem16a* cKO mice were represented by a single focus, reminiscent of frequency responses in control A1 lower than 24 kHz (Fig. 7D). Similarly, at P15-P16, the tone-responsive area of A1 at 80 dB SPL evoked by range of frequencies (3-48 kHz) was reduced (Fig. 7F, G) and the cumulative tone responsive area of auditory cortex was smaller in *Tmem16a* cKO mice (Control, 3.19 mm^2^; cKO, 2.77 mm^2^, Fig. 7H). Multiple high frequency foci were induced in A1 by a 48 kHz tone in older *Tmem16a* cKO mice, but the distances between the centroids of these foci and between low and high frequency foci within A1 were decreased (Fig. 7I, J), indicating macroscopic compression of sound responsive areas in auditory cortex. Together, these results indicate that disruption of early patterned activity ultimately reduces the amount of the brain devoted to processing acoustic information.

**Figure 7.**
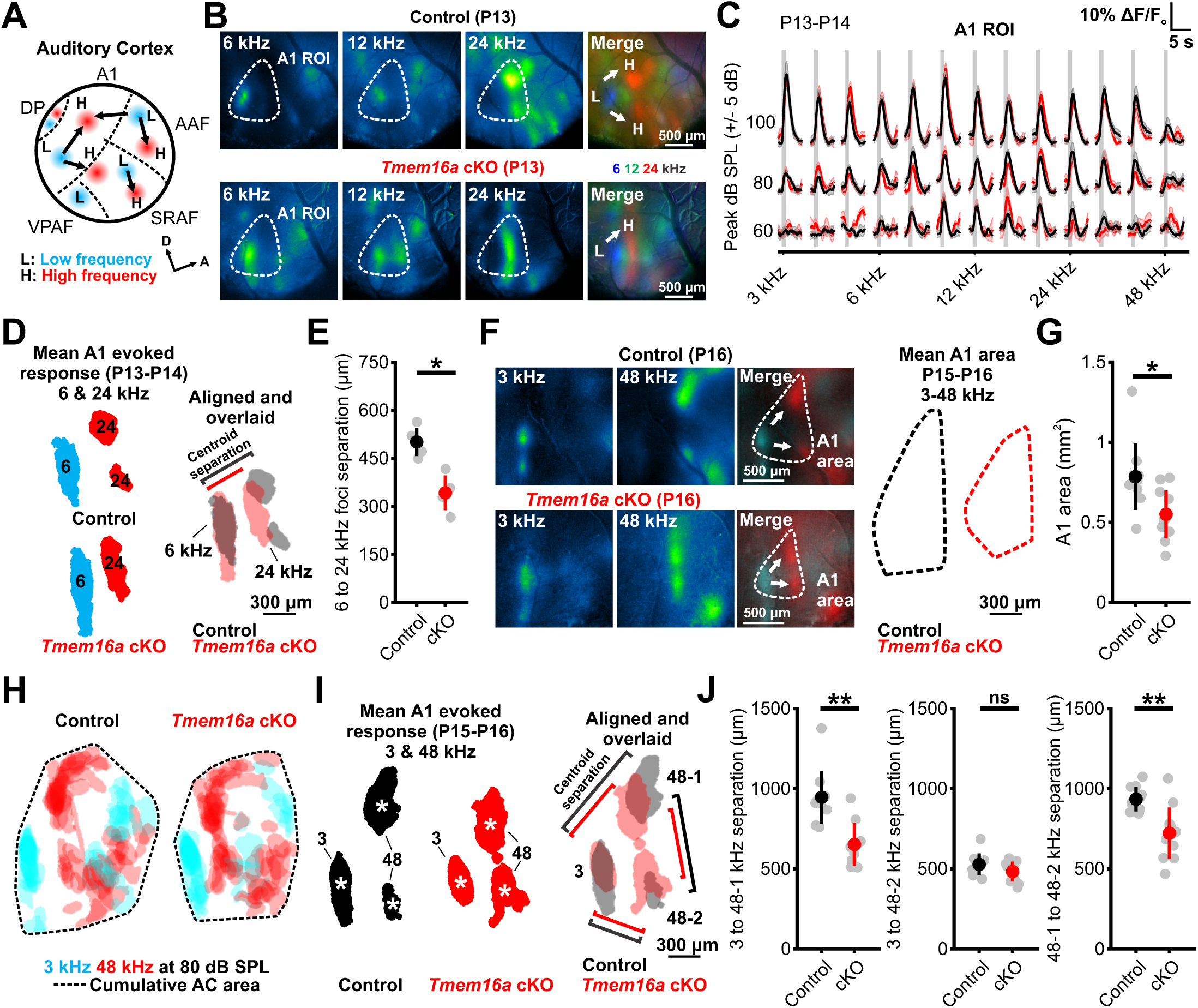
Disruption of pre-hearing spontaneous activity elicits spatial compaction of auditory cortex. (A) Schematic depicting tonotopic organization of mouse auditory cortex, adapted from (Issa *et al*., 2014; Romero *et al*., 2020). L: low frequency, H: high frequency, A1: primary auditory cortex, AAF: anterior auditory field, SRAF: suprarhinal auditory field, VPAF: ventral posterior auditory field, DP: dorsal posterior. (B) Tone-evoked neural calcium transients in AC from P13 control (*Tmem16a^fl/fl^;Snap25-T2A-GCaMP6s*) and P14 *Tmem16a* cKO (*Tecta-Cre;Tmem16a^fl/fl^;Snap25-T2A-GCaMP6s*) mice at 80 dB SPL. Merged image shows tonotopic segregation of pseudocolored pure tone responses along tonotopic axes. (C) Quantification of tone-evoked fluorescence in A1 across a range of frequency and sound level stimuli in P13-P14 control and *Tmem16a* cKO mice. Vertical gray bar indicates tone presentation. n = 6 control mice, 6 *Tmem16a* cKO mice, mean ± SEM. (D) (left) Mean A1 evoked responses to 6 kHz (cyan) and 24 kHz (red) pure tones at 80 dB SPL in P13-P14 control and *Tmem16a* cKO mice. (right) Mean 6 and 24 kHz pure tone responses from control and *Tmem16a* cKO mice aligned to the 6 kHz centroid and overlaid. Lines indicate mean centroid separation between low and high frequency foci. (E) Quantification of centroid separation between 6 and 24 kHz foci. n = 4 control, 4 *Tmem16a* cKO mice; p = 0.0286, Wilcoxon rank sum test. (F) (left) Tone-evoked neural calcium transients in AC from a P16 control and *Tmem16a* cKO mouse at 80 dB SPL. Merged image shows lower (3 kHz) and upper (48 kHz) bound locations of a frequency range in A1. (right) Mean A1 tone-responsive area activated by a 3-48 kHz frequency range. n = 10 control mice, 11 *Tmem16a* cKO mice, (G) Quantification of mean A1 area. n = 10 control mice, 11 *Tmem16a* cKO mice; p = 0.0124, Wilcoxon rank sum test. (H) Cumulative tone responses in auditory cortex of P15-P16 mice to 3 kHz and 48 kHz stimuli at 80 dB SPL. Black dashed line indicates cumulative tone response area. n = 9 control, 10 *Tmem16a* cKO mice. (I) (left) Mean A1 evoked responses to 3 kHz and 48 kHz pure tones at 80 dB SPL in P15-P16 control and *Tmem16a* cKO mice. Centroids of pure tone foci are indicated by white asterisks. (right) Mean 3 and 48 kHz pure tone responses from control and *Tmem16a* cKO mice aligned to the 3 kHz centroid and overlaid. Lines indicate mean centroid separation between low and high frequency foci. (J) Quantification of centroid separation between low (3) and high (48-1 and 48-2) frequency foci in A1. n = 9 control, 10 *Tmem16a* cKO mice; p = 0.0044, 0.2875, 0.0065, Wilcoxon rank sum test.

## Discussion

Neurons in nascent sensory processing networks fire periodic bursts of action potentials prior to the onset of sensory experience. *In vivo* measures of spontaneous activity (Sonntag *et al*., 2009; Clause *et al*., 2014; Babola *et al*., 2018) indicate that each auditory neuron will experience more than 30,000 discrete bursts (∼2.0 bursts/minute; ∼2900 bursts/day) prior to the onset of hearing. This correlated activity propagates from the cochlea to the auditory cortex, mimicking aspects of stimulus encoding that may enable networks to process future sensory input (Kirkby *et al*., 2013; Ge *et al*., 2021; Tiriac *et al*., 2022). Assessing the impact of this prominent, intrinsically generated activity on the functional organization of the auditory system has been difficult, as permanent manipulations of the auditory system necessarily disrupt both spontaneous activity and later processing of sound. Complete silencing of auditory nerve activity during the prehearing period through cochlear ablation or pharmacological inhibition results in dramatic circuit reorganization; however, these manipulations are accompanied by loss of trophic support and substantial neuron loss (Tierney, Russell and Moore, 1997; Mostafapour *et al*., 2000). Similarly, models of congenital deafness exhibit impaired synapse pruning, reduced axonal refinement, and disrupted tonotopic ion channel gradients in auditory brainstem before hearing onset (Leake *et al*., 2006; Leao *et al*., 2006; Hirtz *et al*., 2011; Müller *et al*., 2019), but later deafness of these animals limits functional interrogation of auditory centers. More subtle manipulations of spontaneous activity patterns through genetic alteration of efferent feedback to the cochlea impaired refinement and maturation of MNTB-LSO synaptic connectivity, disrupted the tonotopic gradient of electrophysiologic properties of neurons in the MNTB, and increased central auditory thresholds (Clause *et al*., 2014; Di Guilmi *et al*., 2019; Wang *et al*., 2021); however, these mice retain dysfunctional olivocochlear feedback after hearing onset. Here, we leveraged new insight into the molecular mechanisms that initiate burst firing in the developing cochlea to selectively disrupted this highly stereotyped activity within auditory centers prior to hearing onset, while preserving the ability of the cochlea to transduce sound after hearing begins. We show that this developmental patterned activity is necessary to establish proper acoustic sensitivity, sharpen the frequency tuning of auditory neurons and form future auditory responsive domains in the midbrain and cortex. These results indicate that subtle disruption of activity patterns prior to hearing onset, with no detectable cell loss, is sufficient to induce profound developmental changes in sound processing networks.

### Cochlear supporting cells induce bursts of neural activity in the developing auditory system

In the pre-hearing cochlea, supporting cells within Kölliker’s organ initiate spontaneous neural activity through release of ATP, engagement of purinergic P2RY1 receptors, and activation of TMEM16A. Pharmacologic or genetic disruption of this mechanism alters patterns of spontaneous activity in IHCs, SGNs, and neurons of the MNTB and IC, reducing burst frequency (Wang *et al*., 2015; Babola *et al*., 2020; Maul *et al*., 2022). However, silencing supporting cell activity reduced, but did not eliminate the spontaneous activity of IHCs and SGNs or block correlated firing of central auditory neurons prior to hearing onset, suggesting that there are other mechanisms that may contribute to burst firing at this age. Our prior studies indicate that loss of TMEM16A or P2RY1 causes a ∼10 mV depolarizing shift in the membrane potential of IHCs, due to buildup of extracellular potassium when extracellular space around these cells collapses (Wang *et al*., 2015; Babola *et al*., 2020). As a result, IHC membrane potential is near threshold, allowing these cells to generate calcium spikes with minimal stimulus, leading to spurious, spatially uncoordinated activity. This emergent IHC activity may engage central auditory neurons, but without the precise, spatially restricted coordination afforded by supporting cell-mediated excitation. This residual IHC activity may also be enhanced by direct activation of ionotropic purinergic receptors on IHCs (e.g. P2RX2) (Wang *et al*., 2015; Kolla *et al*., 2020). In this scenario, IHC activity would be expected to be spatially and temporally correlated with ISC activity; however, the correlation between ISC and IHC activity was greatly reduced in *Tmem16a* cKO mice (Supplementary Fig. 3E-H), suggesting that there is minimal residual contribution of purinergic signaling in these animals. When deprived of synaptic input, SGNs become hyperexcitable and respond directly to ISC potassium transients (Babola *et al*., 2018). Similar enhanced excitability may occur in neurons throughout the auditory pathway, progressively amplifying weak inputs and predisposing these circuits to paroxysmal activity, manifesting in the IC-wide events observed in *Tmem16a* cKO mice.

### Gain control in developing sensory pathways

Features of acoustic stimuli, such as sound intensity (loudness) and pitch (frequency), convey critical information needed for localization and interpretation of sounds used for environmental awareness and vocal communication. Our studies in awake mice reveal that when deprived of prehearing burst firing, the gain of sound evoked responses in auditory midbrain was abnormally elevated, with higher amplitude neural responses and broader isofrequency lamina particularly evident to suprathreshold stimuli. These results suggest that nascent sound processing networks are designed to maximize initial responsiveness, in accordance with the overproduction of neurons and excitatory synaptic connections, higher intrinsic excitability of neurons, and delayed maturation of inhibition in the developing brain (Scott, Mathews and Golding, 2005; Dorrn *et al*., 2010; Sun *et al*., 2010; Wong and Marín, 2019). Early patterned activity is then used to balance excitation and inhibition within these networks to enable proper responsiveness and prevent runaway excitation at hearing onset (Vale and Sanes, 2000, 2002; Kotak, Takesian and Sanes, 2008). Similar adaptation occurs in the mature auditory system, in which attenuation of peripheral input and hearing loss triggers gain enhancement necessary to preserve acoustic detection thresholds (Kotak *et al*., 2005; Auerbach, Rodrigues and Salvi, 2014; Chambers *et al*., 2016).

While prominent in subcortical regions, neuronal responsiveness was not significantly increased in AC, suggesting that central auditory centers rely on distinct mechanisms to refine response properties. Notably, while burst firing within the IC is abolished by cochlear removal at P7 (Tritsch *et al*., 2010; Babola *et al*., 2018), sound processing centers in AC receive additional excitatory drive from local cortical activity and cross-modal inputs (Siegel *et al*., 2012; Gribizis *et al*., 2019), which may enable homeostatic reduction in excitability when deprived of peripheral auditory input. As sound evoked neuronal calcium responses in AC were comparable between control and *Tmem16a* cKO mice across the frequency spectrum at hearing onset (Fig. 7C), these results suggest that burst firing triggers global shifts in gain across multiple discrete inputs within the cortex, consistent with current models of homeostatic scaling that manifest through global AMPA receptor internalization, synapse loss, and decreases in intrinsic excitability (Turrigiano, 2012).

The magnitude of gain enhancement in IC of *Tmem16a* cKO mice was highest in regions with greatest acuity, and gradually declined with decreasing stimulus frequency. Auditory circuits adapted for processing higher frequencies appear prone to maladaptive gain enhancement, as tinnitus primarily manifests as a perception of high pitch ‘ringing’ and hyperacusis is most acute at higher frequencies (Stouffer and Tyler, 1990; Gu *et al*., 2010; Hébert, Fournier and Noreña, 2013). While likely important for maintaining activity levels critical for neural survival and circuit refinement, this increased gain in developing sensory pathways could prove detrimental for normal circuit function in maturity and may contribute to developmental disorders in which sensory hypersensitivity is common (Penagarikano, Mulle and Warren, 2007; Marco *et al*., 2011).

### Refinement of sensory receptive fields by intrinsic neural activity

Sensory neurons exhibit precise tuning to specific environmental features. This sensitivity emerges initially through axon guidance cues that establish coarse organization, followed by neural activity-dependent refinement via synapse strengthening or elimination (Katz and Shatz, 1996; Huberman, Feller and Chapman, 2008; Sitko and Goodrich, 2021). In the visual, olfactory, and somatosensory systems, early spontaneous activity has been shown to play crucial roles in refinement of sensory neuron projections and formation of receptive fields (Grubb *et al*., 2003; Yu *et al*., 2004; Chandrasekaran *et al*., 2005; Mrsic-Flogel *et al*., 2005; Burbridge *et al*., 2014; Che *et al*., 2018; Mizuno *et al*., 2018; Antón-Bolaños *et al*., 2019). Like the topographic organization of other sensory pathways, tonotopic information is processed within discrete domains throughout the auditory system, enabling isolation of frequency components critical for differentiating pitch and deconvolving complex sounds. However, assessment of functional evoked responses *in vivo* at both global and cellular levels following disruption of correlated pre-hearing spontaneous activity had not yet been previously performed. In *Tmem16a* cKO mice, individual IC neurons displayed broader tuning, and correspondingly, the spatial representation of pure tones across the tonotopic axis in IC was expanded, indicating that early patterned neural activity initiates functional refinement within auditory processing centers. However, it is also possible that the infrequent, large-scale events observed in *Tmem16a* cKO mice may contribute to the expansion of frequency tuning by inducing inappropriate coordinated firing of neurons across tonotopic domains through Hebbian mechanisms. Nevertheless, our results suggest that precise, coordinated firing of hair cells and neurons within discrete domains during early development is necessary to achieve proper tuning and frequency partitioning.

A recent study by Maul et al. (2022) used a similar approach to explore the role of spontaneous activity in auditory development, revealing that genetic inactivation of *Tmem16a* resulted in broader neural tuning of neurons in the MNTB and impaired refinement of MNTB-LSO functional connectivity, consistent with our observations in the IC. However, MNTB neurons from *Tmem16a* cKO mice demonstrated reduced firing rates at their characteristic frequency and a ∼5 dB increase in thresholds, suggestive of reduced acoustic sensitivity to low intensity stimuli in the MNTB. Notably, we observed similar subtle trends in single cell threshold and calcium evoked amplitude to low frequency, low intensity stimuli in *Tmem16a* cKO mice (Supplementary Fig. 10A-C), but amplitudes dramatically increased at suprathreshold sound levels. The use of ketamine/xylazine anesthesia by Maul et al., which has been shown to alter spike rate, timing, and threshold in auditory neurons (Jing, Pecka and Grothe, 2021) and disrupt neuromodulatory transmission, may account for this difference. Additionally, the authors relied on *Pax2-Cre* to remove *Tmem16a*, a line that displays both incomplete knockout within the inner ear (Wang *et al*., 2015; Eckrich *et al*., 2019) and extensive recombination in the CNS (Ohyama and Groves, 2004), resulting in a hypomorph that retains spontaneous activity in ISCs (Wang *et al*., 2015), with unknown consequences.

### Neuronal burst firing establishes central auditory processing domains

Sensory domains in the cerebral cortex are shaped through competitive interactions, and loss of peripheral input results in atrophy of neural territories devoted to processing that input (Rauschecker *et al*., 1992; Kahn and Krubitzer, 2002; De Villers-Sidani *et al*., 2007; Moreno-Juan *et al*., 2017). Disruption of pre-hearing spontaneous activity led to spatial compaction of auditory domains, reducing the areas within IC and AC devoted to processing tonal sound information. These findings indicate that distinct patterns of activity, not simply the presence of neural activity, provides important cues to define future organization and functionality. Whether these topographic changes arise from competition with other sensory modalities, or simply reflect activity-dependent stabilization, is not yet known. Moreover, it remains to be determined whether normal acoustic input can reverse this topographic compression of auditory domains, comparable to what can be achieved in the visual system in patients with amblyopia (Clarke *et al*., 2003). Our results suggest that genetic variants associated with deafness which disrupt central spontaneous activity patterns (Babola *et al*., 2018; Shrestha *et al*., 2018; Sun *et al*., 2018; Müller *et al*., 2019), may induce profound changes in the functional organization of auditory centers. Understanding the distinct changes induced by these different activity patterns may reveal new ways to enhance the performance of prosthetic hearing devices for individual patients suffering from congenital hearing loss.

## Acknowledgements

We thank members of the Bergles laboratory for discussions and comments on the manuscript. We thank Michele Pucak and Abigail Bush in the Multiphoton Imaging Core and Terry Shelley in the Neuroscience Machine Shop for assistance. Funding was provided by grants from the National Institutes of Health (DC008060, NS050274) to Dwight E. Bergles. Calvin J. Kersbergen is supported by an individual NRSA fellowship (DC018711) and a Medical Scientist Training Program training grant (GM136577) from the National Institutes of Health.

## Author Contributions

Calvin J. Kersbergen, Conceptualization, Methodology, Investigation, Formal analysis, Funding acquisition, Writing--original draft; Travis A. Babola, Methodology, Investigation, Formal analysis, Writing--review and editing; Jason Rock, Tools and reagents, Writing--review and editing; Dwight E. Bergles, Conceptualization, Methodology, Supervision, Funding acquisition, Writing--original draft, Writing--review and editing.

## Competing Interests

Dwight E. Bergles is a paid consultant of Decibel Therapeutics.

Correspondence should be addressed to Dwight E. Bergles at

## STAR Methods

### RESOURCE AVAILABILITY

#### Lead contact

Requests for sharing resources, tools, code, and reagents should be directed to the corresponding author Dwight E. Bergles.

#### Materials, data, and code availability

This study did not generate any unique tools or reagents. Code used for data analysis and figure generation is deposited on Github (https://github.com/ckersbe1/Cochlea-spont-activity-development-MS).

### EXPERIMENTAL MODEL AND SUBJECT DETAILS

#### Animals

This study was performed in strict accordance with the recommendations provided in the Guide for the Care and Use of Laboratory Animals of the National Institutes of Health. All experiments and procedures were approved by the Johns Hopkins Institutional Care and Use Committee (Protocols #M018M330, M021M290). Generation and genotyping of transgenic mice (*Tmem16a* floxed mice (Schreiber *et al*., 2015), *Tmem16a-GFP* mice (Huang *et al*., 2012), *Tecta-Cre* mice (JAX Stock No. 035552) (Babola *et al*., 2020)*, R26-lsl-eGFP* mice (MMRRC Stock No. 32037) (Sousa *et al*., 2009), *R26-lsl-GCaMP3* mice (Jax Stock No. 028764) (Paukert *et al*., 2014), and *Snap25-T2A-GCaMP6s* mice (JAX Stock No. 025111) (Madisen *et al*., 2015)) have been previously described. Mice were maintained on a mixed C57Bl/6NJ-FVB/NJ background. Both male and female mice were used for all experiments in equal numbers at ages P1-P21 (precise age for each experiment can be found in the figure legends). Breeding pairs were checked daily in the morning for pups, with the date of first observation of pups defined as P0. Mice were housed on a 12-hour light/dark cycle and were provided food ad libitum.

### METHOD DETAILS

#### Electrophysiology

Apical segments of the cochlea were acutely isolated from P0-P12 mouse pups and used within 2 hours, as described previously (Babola *et al*., 2021). Cochlea were superfused by gravity at ∼2 mL/minute with bicarbonate-buffered aCSF at physiologic temperature (32-34 C) containing (in mM): 119 or 115 NaCl, 2.5 or 6 KCl, 1.3 MgCl_2_, 1.3 CaCl_2_, 1 NaH_2_PO_4_, 26.2 NaHCO_3_, 11 D-glucose saturated with 95% O_2_ / 5% CO_2_ at a pH of 7.4. Whole cell recordings from ISCs and IHCs were made under visual guidance using differential interference contrast (DIC) transmitted light using a 40x magnification objective on a Zeiss Axioskop 2 microscope. Electrodes had tip resistances between 2.5 and 4.0 MΩ (ISCs) or between 4.0 and 6.0 MΩ (IHCs) with internal solution of (in mM): 134 KCh_3_SO_3_, 20 HEPES, 10 EGTA, 1 MgCl_2_, 0.2 Na-GTP, pH 7.4. Measurements of membrane resistance in ISCs were obtained immediately through repeated 10 mV voltage steps from -20 to +20 mV relative to the holding potential. Spontaneous currents were recorded from ISCs held at -90 mV (close to resting potential) for a minimum of 10 minutes. Spontaneous currents were recorded with IHCs held at -70 mV (close to resting potential) for a minimum of 10 minutes. Recordings were performed using pClamp 9 software with a Multiclamp 700A amplifier (Axon Instruments), low pass Bessel filtered at 1 kHz, and digitized at 5 kHz (Digidata 1322a, Axon Instruments). Recordings exhibiting > 20% change in access resistance or with access resistance > 30 MΩ at the start of recording were discarded. Errors due to voltage drop across the series resistance and liquid junctional potential were left uncompensated. Analysis of input resistance and spontaneous activity was performed offline in MATLAB (Mathworks). Spontaneous currents were detected using the ‘peakfinder’ function, with a fixed peak threshold (baseline + 3 standard deviations) and minimum peak amplitude (10 pA for ISCs, 5 pA for IHCs).

#### Transmitted light imaging

For time-lapse imaging of spontaneous osmotic crenations in supporting cells, acutely excised cochleae were visualized using DIC optics through a 40x water-immersion objective coupled to a 1.8x adjustable zoom lens as described above. Images were acquired at one frame per second using a frame grabber (LG-3; Scion) and Scion Image software. Crenations were detected by generation of difference movies in MATLAB through subtraction of frames at time t_n_ and t_n+5_ seconds, as described previously (Babola *et al*., 2020). Filled areas represent temporally pseudo-colored detected events overlaid on the transmittance image.

#### Cochlea Explant Culture and Calcium Imaging

Cochlea segments were acutely isolated from P6-P7 mice in ice-cold, sterile filtered, HEPES buffered artificial cerebrospinal fluid (aCSF) containing (in mM): 130 NaCl, 2.5 KCl, 10 HEPES, 1 NaH_2_PO_4_, 1.3 MgCl_2_, 2.5 CaCl_2_, 11 D-glucose, as previously described (Zhang-Hooks *et al*., 2016; Babola *et al*., 2018). Explants were mounted onto Cell-Tak (Corning) treated coverslips and incubated at 37 C for 12 hours in Dulbecco’s modified Eagle’s medium (F-12/DMEM; Invitrogen) supplemented with 1% fetal bovine serum (FBS) and 10 U/mL penicillin (Sigma) prior to imaging. After overnight culture, cochleae were transferred to the recording chamber and superfused with bicarbonate-buffered aCSF at physiologic temperature (32-34 C) containing (in mM): 115 NaCl, 6 KCl, 1.3 MgCl_2_, 1.3 CaCl_2_, 1 NaH_2_PO_4_, 26.2 NaHCO_3_, 11 D-glucose saturated with 95% O_2_ / 5% CO_2_ at a pH of 7.4. Cochleae were illuminated with a 488 nm laser (maximum 25 mW power), and optical sections containing both IHCs and ISCs were obtained with a pinhole set to 3.67 Airy units, corresponding to 5.4 μm of z-depth. Images were captured at 2 Hz using a using a Zeiss laser scanning confocal microscope (LSM 710) through a 20X objective (Plan APOCHROMAT 20x/1.0 NA) at 512 x 512 pixels (425.1 by 425.1 microns) for a minimum of 10 minutes. Analysis of supporting cell and hair cell calcium transients were performed as previously described (Babola *et al*., 2021). Briefly, images were normalized to ΔF/F_o_ values at the 10^th^ percentile, and a grid of 10 x 10 pixel squares were overlaid on the ISC region, while individual circular ROIs were placed at the basal pole of IHCs.

#### *In vivo* imaging of spontaneous activity

Installation of neonatal cranial windows has been previously described (Babola *et al*., 2018). Briefly, mice were anesthetized in inhaled isoflurane (4% induction, 1.5% maintenance), the dorsal skull exposed to allow for headbar implantation, and a cranial window placed over the resected intraparietal bone overlying the midbrain. After >1 hour of post-surgical recovery from anesthesia, neonatal mice were moved into a 15 mL conical tube and head-fixed under the imaging microscope. During imaging, pups were maintained at 37 C using a heating pad and temperature controller (TC-1000; CWE). Wide field epifluorescence images were captured at 10 Hz using a Hamamatsu ORCA-Flash4.0 LT digital CMOS camera attached to a Zeiss Axio Zoom.V16 stereo zoom microscope at 17x magnification illuminated continuously with a metal halide lamp (Zeiss Illuminator HXP 200C). Each recording of spontaneous activity consisted of uninterrupted acquisition over a minimum of 10 minutes. Two photon imaging was performed using a Zeiss 710 LSM microscope with two-photon excitation achieved by a Ti:sapphire laser (Chameleon Ultra II; Coherent) tuned to 920 nm. Images were collected at 4 Hz (256 x 256 pixels, 425 x 425 μm) from 150 μm Z-depth in the central IC for a minimum of 10 minutes.

Widefield imaging analysis followed previously described methods in MATLAB (Babola *et al*., 2018). Image intensities were normalized as ΔF/F_o_ values, where ΔF = F – F_o_ and F_o_ was defined as the 10^th^ percentile value for each pixel. Oval regions of interest were placed over the right and left IC, and signal peaks were identified using built-in peak detection (‘findpeaks’) with a fixed threshold (2% ΔF/F_o_) and minimum peak amplitude (1% ΔF/F_o_). For analysis of spatial band width, a 25 x 100 rectangular ROI rotated 45-55 degrees was placed over each inferior colliculus aligned with the tonotopic axis of the IC (Babola *et al*., 2018). The rectangle was averaged along the short axis, creating a 100 x 1 line scan of the tonotopic axis of the IC for the duration of the time series. Events were detected using the function ‘imregionmax’ with a fixed threshold (2% ΔF/F_o_). Line scans of individual events were normalized to the maximum ΔF/F_o_ at that time point, and the band width calculated as the length along the tonotopic axis above the 75^th^ percentile of the peak ΔF/F_o_. Two-photon imaging analysis was performed as previously described (Kellner *et al*., 2021). Briefly, an array of 10 x 10 pixel grid ROIs was placed over the normalized ΔF/F_o_ image, and individual ROIs were considered responsive when the signal exceeded the median + 2 standard deviations. Coordinated events were defined as simultaneous activation of >2 adjoining ROIs for > 1.5 s.

#### Analysis of neuronal activity in the superior colliculus

For assessment of spontaneous neuronal activity in the superior colliculus driven by retinal ganglion cell burst firing (Ackman, Burbridge and Crair, 2012), images were normalized to ΔF/F_o_ values at the 10^th^ percentile, as described above. 200 x 150 pixel ROIs were placed over each colliculi. Pixels within each ROI were downsampled by a factor of 5 and considered active if they exceeded the mean + 3 standard deviations for that pixel. Retinal waves were defined as periods > 1 s where > 5 pixels were simultaneously active. Wave duration was defined as the total time in which > 5 pixels were continuously active during a given retinal wave.

#### Cochlea and brain immunohistochemistry

Mice were deeply anesthetized with intraperitoneal injection of 10 mg/ml sodium pentobarbital (>P14) or isoflurane overdose (<P10) and cardiac perfused with ice cold 1X PBS followed by 4% paraformaldehyde (PFA) in 0.1 M phosphate buffer, pH 7.4. The brain and inner ears were carefully removed from the skull, post-fixed in 4% PFA overnight at 4 C, and stored in 1X PBS with 0.1% sodium azide. For cross sections, cochleae were decalcified in 10% EDTA in 0.1 M phosphate buffer (pH 7.4) at 4 C (P7: 2-4 hours, >P14: 48 hours), cryopreserved in 30% sucrose, embedded in O.C.T. compound (Tissue Tek), cut in 10 μm sections using a cryostat, and placed directly on slides (SuperFrost Plus, Fisher). For immunostaining, cochlea sections and free-floating whole mount cochlea were preincubated in blocking solution (0.3-0.5% Triton X-100, 5% Normal Donkey Serum in PBS, pH 7.4) and incubated overnight at 4 C (cochlea) or room temperature (brain) with primary antibodies (Chicken anti-GFP, 1:4000, Aves; Rabbit anti-MyosinVIIA, 1:300, Proteus Biosciences; Rabbit anti-β3-tubulin, 1:500, Cell Signaling Technologies). Following overnight incubation, tissues were washed 3 x 10 minutes in PBS and incubated with corresponding donkey secondary antibodies (Alexa Fluor 488, Alexa Fluor 546, or Alexa Fluor 647; 1:2000, Invitrogen) for 2-3 hours at room temperature. Finally, tissue was washed 3 x 10 minutes in PBS, incubated with 1:10000 DAPI in PBS, and sealed using Aqua Polymount (Polysciences, Inc). For hematoxylin and eosin staining, cochleae were decalcified and dehydrated in 70% ethanol, embedded in paraffin, cut in 5 μm sections, and stained by the Reference Histology core in the Department of Pathology at Johns Hopkins Hospital. Images were captured using an epifluorescence and light microscope (Keyence BZ-X) or a laser scanning confocal microscope (LSM 880, Zeiss). For analysis of hair cell density, decalcified cochleae were dissected and cut into 3 segments of equal length (apex, middle, base), and stained for hair cell markers as described above. Images were collected at 25x magnification at the mid-portions of each segment corresponding roughly to 8 kHz, 24 kHz, and 50 kHz, respectively. For analysis of spiral ganglion neurons, 10 μm-thick mid-modiolar cross-sections were identified, and images collected from the apical, middle, and basal spiral ganglia visible within that section. SGN density was determined in a blinded manner by measuring spiral ganglion area in ImageJ and subsequent manual counts of all visible Tuj1-labeled neuronal soma. If mid-modiolar sections were visualized across multiple sections and slides, images were collected from all regions and averaged.

#### Auditory brainstem response measurements

For assessments of auditory brainstem responses (ABRs), mice were anesthetized by intraperitoneal injection of Ketamine (100 mg/kg) and Xylazine (20 mg/kg) and placed in a sound attenuation chamber. Body temperature was maintained at 37 C using an isothermal heating pad. Subdermal platinum needle electrodes (E2, Grass Technologies) were placed posterior to the pinna, at the vertex, and leg (ground). Acoustic stimuli consisted of 0.1 ms clicks and 5 ms tone pips (2 ms rise time) of varying frequency (8, 16, 24, and 32 kHz) presented at a rate of 40 Hz. Stimuli were generated by a RZ6 processer (Tucker Davis Technologies) at sound pressure levels of 90-20 dB in 5 dB descending increments and delivered through a free-field speaker (MF-1, Tucker Davis Technologies) placed 10 cm away from the pinna. Calibration of the free-field speaker was performed using an ACO Pacific microphone (7017) and preamplifier (4016). ABR signals were amplified (Medusa 4Z, Tucker Davis Technologies), band-pass filtered (300 Hz and 3 kHz), digitized (RZ6 Processor, Tucker Davis Technologies), and averaged across 600-700 stimuli. ABR thresholds were calculated in an automated manner offline in MATLAB as the lowest stimulus intensity determined by linear interpolation that produced peak-to-peak ABR signals that were greater than 2 standard deviations above the peak-to-peak background signal. P1-N1 amplitudes and peak timing were determined using the ‘findpeaks’ function with fixed criteria. Displayed ABR traces represent mean +/- standard deviation (shaded region) across all animals.

#### Sound-evoked calcium imaging

Cranial windows over inferior colliculus in P13-P15 mice were performed as previously described for P7 animals, with animals allowed to recover a minimum of 2 hours prior to imaging. To enable image registration for two-photon imaging, aCSF containing Sulfarhodamine 101 (10 μM) was washed over the brain surface before sealing of the cranial window to label astrocytes (Nimmerjahn and Helmchen, 2012). Auditory cortex (AC) windows were installed using a microblade and microscissors to remove a ∼4 mm circular region of skull overlying the right auditory cortex. Following skull removal, the dura was carefully removed using microscissors and exposed brain was continuously immersed in aCSF until a 5 mm coverslip was attached using superglue. During imaging, awake animals were head-fixed but on a freely rotating tennis ball to simulate natural movement. Wide field epifluorescence images were captured as described above, with 21x magnification for AC imaging. Two photon images were captured as described above, with some modifications. Low magnification mapping was performed in the central nucleus of the IC over a 425 x 425 μm field of view at 150-200 μm Z-depth at 2 Hz (512 x 512 pixels). Following coarse mapping of pure tone responses in the IC, high magnification images were obtained along the tonotopic axis (centered on the 6 kHz response location) to assess of single cell tuning and gain through a 212 x 212 μm field of view continuously visualized at 5 Hz (256 x 256 pixels).

Acoustic stimuli were generated within the RPvdsEx software (Tucker Davis Technologies), triggered using the microscope’s frame out (widefield) or line out (two-photon) signal, and delivered through the RZ6 Processor (Tucker-Davis Technologies). Stimuli were presented using a free-field speaker (MF-1, Tucker Davis Technologies) placed 10 cm from the left ear within a custom sound attenuation chamber with external noise attenuation of 40 dB (Babola *et al*., 2018). Calibration of the free-field speaker was performed using an ACO Pacific microphone (7017) and preamplifier (4016). Given the flat intensity profile (peak 100 dB SPL +/- 5 dB for a 2.0 V stimulus across all tested frequencies), levels were not corrected across presented frequencies. For widefield imaging, stimuli consisted of 4 repetitions of sinusoidal amplitude modulated (SAM) pure tones (1 s, 10 Hz modulation) from 3 to 48 kHz in ¼ octave intervals. All stimuli were cosine-squared gated (5 ms) and played in a random order at 5 s intervals. Stimuli were presented from 100 dB to 40 dB SPL in 10-20 dB attenuation steps. For two-photon low magnification imaging, acoustic stimuli consisted of 1 s SAM tones from 3-24 kHz at 90 dB SPL. For two photon high magnification imaging, acoustic stimuli consisted of 200 ms SAM tones presented in a random order ranging from 3 to 24 kHz with 0.5 to 0.25 octave steps at 90-30 dB SPL (5-10 dB attenuation steps) with 5 s inter-stimulus intervals.

For analysis of widefield sound-evoked responses, raw images underwent bleach correction and normalization as described above for widefield imaging of spontaneous activity. Image segments were separated by tone frequency, aligned from 1 second prior to and 3 seconds following tone presentation, and averaged across the 4 presentations of each tone. Normalized and averaged images were used for display purposes. For analysis of evoked amplitude and auditory thresholds in inferior colliculus, an oval ROI was placed over the contralateral IC. For assessment of spatial band width and frequency mapping, a maximum-intensity projection of the mean sound-evoked response for each presented frequency was rotated by 45-55 degrees and a 25x250 pixel rectangle was placed along the tonotopic axis, centered at the 3 kHz response location for each animal. To generate the sound-evoked spatial profile, ΔF/F_o_ signal intensity was averaged along the short rectangle axis. Profiles were normalized to the maximum ΔF/F_o_ intensity (peak response location) along the tonotopic axis.

The normalized spatial band width of the evoked response was defined as the width of the 75^th^ percentile of the normalized response for each mouse. For analysis of auditory cortex response amplitudes and time course, a circular ROI was placed over A1 after visualization of averaged pure tone responses based on previously described mesoscopic maps of auditory cortex in adult mice (Issa *et al*., 2014; Romero *et al*., 2020). The low and high frequency borders of A1 were defined by their caudal and dorsal locations in auditory cortex. Thresholding and segmentation of AC pure tone foci was performed by binarizing maximum evoked responses using a fixed threshold. Images were aligned based on the 3 kHz A1 location and rotated by the angle of a vector between the 3 kHz and 48 kHz A1 foci to allow for direct comparison of tonotopic organization between animals. At P13, this process was the same except 6 and 24 kHz stimuli at 80 dB SPL were used (due to inter-animal variability in responses to the frequency extremes (3 and 48 kHz) at P13). A1 area was calculated as the total area bounded by a binarized maximum projection of activated area through a 3-48 kHz frequency sweep at 80 dB SPL (‘bw’, ‘noholes’). Cumulative auditory cortex area was calculated as the boundary encompassing the maximum aligned projection of all 3-48 kHz tone responses in all mice. Centroid separation was measured as the Euclidean distance between centroids of binarized low and high frequency foci. If motion correction was required of widefield movies, it was performed using the MOCO fast motion correction plugin (ImageJ) (Dubbs, Guevara and Yuste, 2016).

For analysis of single cell responses, images first underwent motion correction using the ImageJ plugin MultiStackReg, using SR101 signal as a fixed landmark. Next, images were aligned by tone presentation and sound level and averaged across 4-6 repetitions. ROIs were manually placed around individual sound-responsive soma and averaged, and ΔF/F_o_ calculated for each frequency and stimulus level, where ΔF = F – F_o_ and F_o_ represents the 20^th^ percentile value for the ROI in the 1 s prior to tone onset across all stimuli. ROIs were considered responsive to a given stimulus if ΔF/F_o_ in the 1 s following stimulus onset exceeded 5 x standard deviation of the ΔF/F_o_ signal 1 s before stimulus onset. Maximum bandwidth was calculated as the maximum range of frequency responses within a single stimulus intensity level, while gaussian bandwidth was calculated by fitting a single term gaussian (‘fit’, ‘gauss1’) to the maximum evoked amplitude across all frequency stimuli within a given stimulus level.

Frequency response area (FRA) was calculated as the integral of the total positive responses to all frequencies and sound levels with constant stimulus parameters (3-24 kHz, 40-90 dB SPL). Best frequency for a cell was defined as the stimulus frequency that elicited the maximum ΔF/F_o_ response at any sound level. Center frequency for a cell was defined as the stimulus frequency that elicited the maximum ΔF/F_o_ response at the lowest sound level that elicited a response. Neuropil analysis was performed by averaging across the entire field following application of a mask over all neuronal soma within the field identified within a maximum projection image.

### QUANTIFICATION AND STATISTICAL ANALYSIS

All statistical testing was performed in MATLAB, with the results of each statistical test available in MATLAB live scripts for each figure, available at: https://github.com/ckersbe1/Cochlea-spont-activity-development-MS. All data is presented as mean +/- standard deviation, unless otherwise noted. For two-group comparisons, datasets were tested for normality using the Lilliefors test (‘lillietest’). If unable to reject the null hypothesis that the dataset is normally distributed using the Lilliefors test, an unpaired (‘ttest2’) two-tailed t-test was used to compare groups, with the addition of assumption of unequal variances (‘Vartype’, ’unqual’) as necessary. If the null hypothesis of normality was rejected, a nonparametric Wilcoxon rank sum (‘ranksum’) test was used for unpaired samples. If multiple comparisons on the same datasets were made, a Benjamini-Hochberg correction of the false discovery rate (‘fdr_BH’) was made to adjust p values to lower the probability of type 1 errors. Comparison of cumulative distributions was performed using a two-sample Kolmogorov-Smirnov test (‘kstest2’). For datasets with multiple comparisons of non-independent samples or with missing data (ie, no stimuli at a given frequency or sound level, or data removed due to movement artifacts), a repeated measures ANOVA (‘ranova’) or a linear mixed-effects model (‘fitlme’) was used, using the reduced maximum likelihood fit method (‘FitMethod’,’reml’) and the Satterthwaite approximation of degrees of freedom. Use of a linear mixed model enabled accounting for data dependency for repeated measurements from the same mouse as a random effect. Sidák post hoc test was used to assess for post hoc comparisons as indicated in figure legends. Adjusted p-values are displayed as follows: * = p < 0.05, ** = p < 0.01, *** = p < 0.001, **** = p < 0.0001, ns = not significant. Details about number of data points and individual statistical tests and p values can be found in the figure legends.

### KEY RESOURCES TABLE

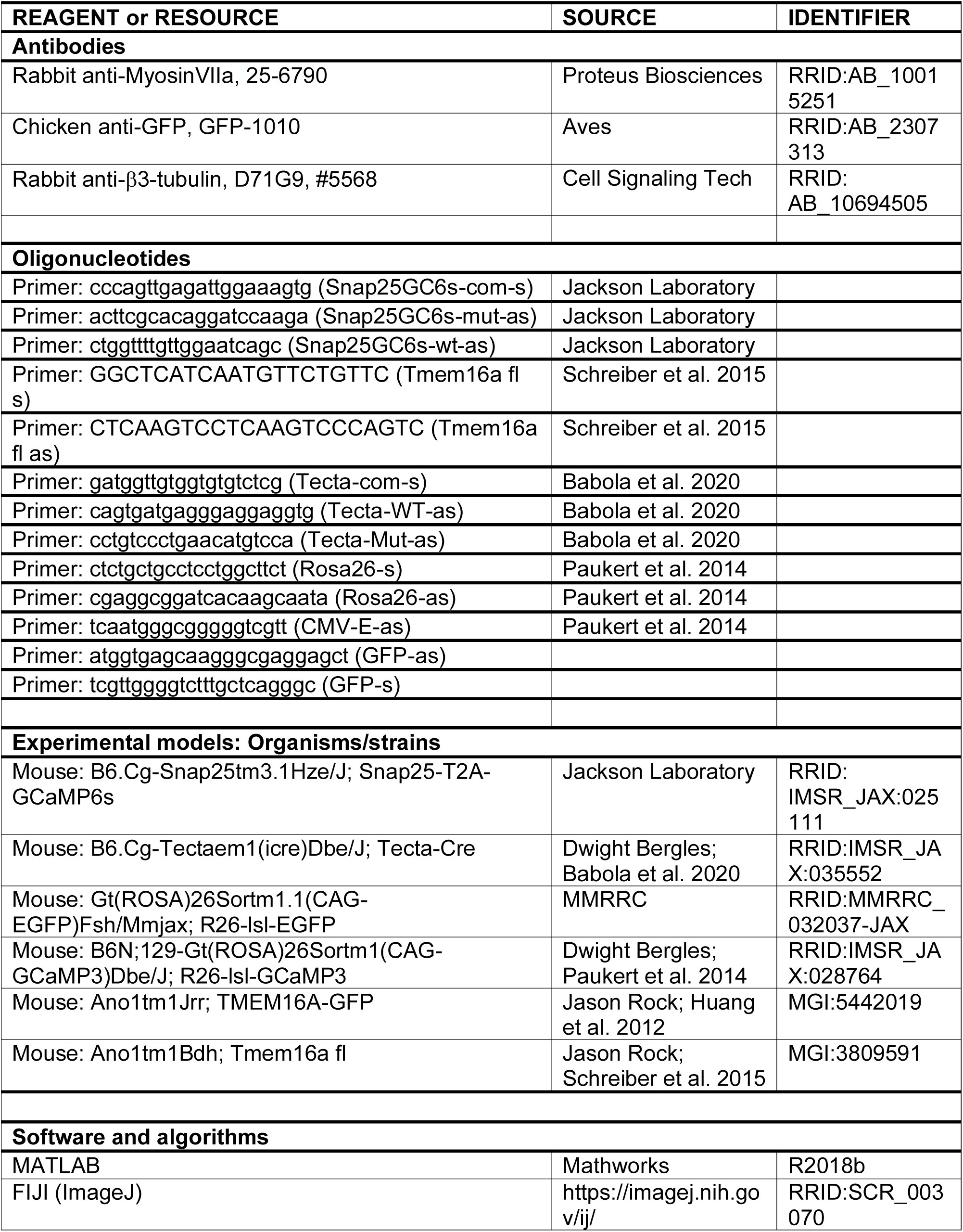
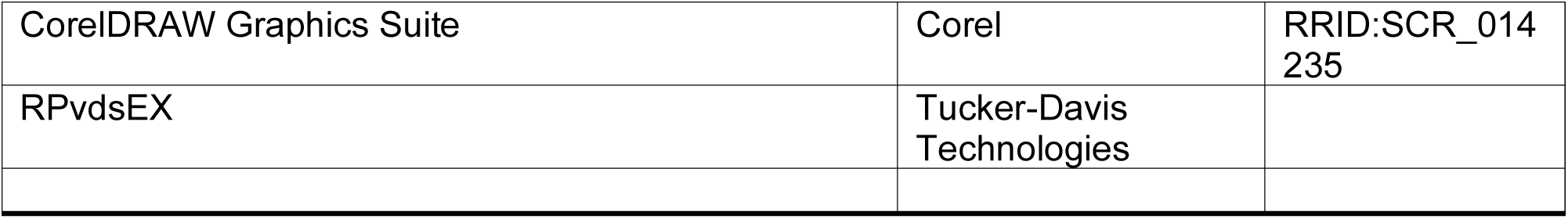

#### Supplementary Material

**Supplementary Figure 1. Recombination of *Tecta-Cre* in the cochlea and CNS, related to** Figure 1

(A) Low magnification image of *Tecta-Cre* recombination, as shown by EGFP (green) reporter expression, in a P7 apical whole mount cochlea. Hair cells are indicated by immunoreactivity to MyoVIIA (magenta).

(B) High magnification of white square in (A) depicting recombination in inner supporting cells (ISCs) in the developing cochlea.

(C) (left) Extent of recombination of *Tecta-Cre* in cross sections of P21 cochlea. White square indicates site of high magnification. (right) Reporter expression is seen within interdental cells, inner sulcus cells, supporting cells of the organ of Corti, and inner and outer hair cells (MyoVIIA, magenta; outlined by black dashed lines).

(D) Extent of recombination of *Tecta-Cre* in sagittal section of P21 brain, as shown by EGFP (white) reporter expression. Recombination is limited within the inferior colliculus (circled). Blue squares indicate sites of high magnification in (F).

(E) Extent of recombination of *Tecta-Cre* in coronal sections of P21 brain, as shown by EGFP (white) reporter expression, in the auditory brainstem, midbrain, thalamus, and cortex.

(F) High magnification of blue squares in (D) depicting sparse recombination in cortical astrocytes and the dentate gyrus.

**Supplementary Figure 2. TMEM16A channels are required for spontaneous activity and osmotic crenation across postnatal development, related to Figure 1**

(A) Whole cell patch clamp recording configuration from inner supporting cells (ISCs).

(B) Whole cell patch clamp recordings of spontaneous activity from P1 control (*Tmem16a^fl/fl^*) and *Tmem16a* cKO (*Tecta-Cre;Tmem16a^fl/fl^*) ISCs within acutely isolated cochleae.

(C) Same as (B), but for P11 cochleae.

(D) Quantification of mean current amplitude in ISCs at three ages (P1, P7, P11) encompassing postnatal development. n = 5, 6, 7 control, 6, 6, 7 *Tmem16a* cKO ISCs (P1, P7, P11); p = 0.0154, 0.0346, 0.0445 (P1, P7, P11), two-sample t-test with unequal variances and Benjamini-Hochberg correction.

(E) Quantification of integral (charge transfer) in ISCs at three ages (P1, P7, P11). n = 5, 6, 7 control ISCs, 6, 6, 7 *Tmem16a* cKO ISCs (P1, P7, P11); p = 0.0154, 0.0452, 0.0061 (P1, P7, P11), Wilcoxon rank sum test (P11) or two-sample t-test with unequal variances and Benjamini-Hochberg correction (P1, P7).

(F) Voltage step protocol and current responses from postnatal day 7 (P7) control and *Tmem16a* cKO ISCs.

(G) Quantification of membrane resistance in ISCs at three ages (P1, P7, P11) encompassing postnatal development. n = 6, 12, 8 control ISCs, 11, 8, 8 *Tmem16a* cKO ISCs (P1, P7, P11); p = 0.3011, 0.4034, 0.6662 (P1, P7, P11), Wilcoxon rank sum test (P1, P11) or two-sample t-test (P7).

(H) Intrinsic optical imaging of osmotic crenations in control and *Tmem16a* cKO cochleae. Detected crenations are indicated with transparent colored areas based on time of occurrence.

(I) Quantification of spontaneous crenation frequency at three ages (P1, P7, P11). n = 5, 7, 6 control cochleae, 5, 6, 6 *Tmem16a* cKO cochleae (P1, P7, P11); p = 0.2199, 0.0012, 0.0022 (P1, P7, P11), Wilcoxon rank sum test (P7, P11) or two-sample t-test (P1).

**Supplementary Figure 3. TMEM16A channels are required for coordinated activation of inner hair cells, related to Figure 1**

(A) Cochlea whole mount calcium imaging paradigm to visualize spontaneous calcium transients within inner supporting cells (ISCs) and inner hair cells (IHCs) using the Cre-dependent genetically encoded calcium indicator GCaMP3. Green depicts coordinated activation of ISCs and IHCs within typical control spontaneous events.

(B) Calcium transients in ISCs, color coded by time of occurrence, in P7 control (*Tecta-Cre;Tmem16a^fl/+^;R26-lsl-GCaMP3*) and *Tmem16a* cKO (*Tecta-Cre;Tmem16a^fl/fl^;R26-lsl-GCaMP3*) cochleae. ISC calcium transients are quantified using a grid-based ROI analysis.

(C) Raster plot of calcium transients within randomly selected ISC grid ROIs.

(D) Quantification of ISC calcium event frequency, mean event amplitude, mean event duration, and mean event area. n = 6 control cochleae, 6 *Tmem16a* cKO cochleae; p = 0.9372, 0.4127, 0.1299, 0.1039 (frequency, amplitude, duration, area), Wilcoxon rank sum test with Benjamini-Hochberg correction.

(E) Maximum intensity projection and fluorescence traces in ISCs and adjacent IHCs, as indicated by ROIs, from a single ISC event in P7 control (left) and *Tmem16a* cKO (right) cochleae. Asterisks indicate two active IHC ROIs during the supporting cell calcium transient that are not temporally aligned with local ISC activity or with each other.

(F) Summed traces of fluorescence in all ISCs (blue), all IHCs (black), and individual IHC ROIs from a P7 control cochlea.

(G) Summed traces of fluorescence in all ISCs (blue), all IHCs (black), and individual IHC ROIs from a P7 *Tmem16a* cKO cochlea.

(H) Quantification of mean correlation (80^th^ percentile) between summed ISC activity and individual IHCs. n = 5 control, 6 *Tmem16a* cKO; p = 0.0043, Wilcoxon rank sum test.

**Supplementary Figure 4. Inner hair cells are depolarized following removal of supporting cell TMEM16A, related to Figure 1.**

(A) Whole cell patch clamp recording configuration from inner hair cells (IHCs).

(B) Quantification of P7 IHC resting membrane potential at P7. n = 5 control IHCs, 4 *Tmem16a* cKO IHCs; p = 0.0317, Wilcoxon rank sum test.

(C) Voltage step protocol and current responses from postnatal day 7 (P7) control and *Tmem16a* cKO IHCs.

(D) Quantification of P7 IHC membrane resistance. n = 5 control IHCs, 4 *Tmem16a* cKO IHCs; p = 0.2857, Wilcoxon rank sum test.

**Supplementary Figure 5. Disruption of auditory spontaneous activity does not alter retinal wave frequency, related to Figure 2**

(A) *In vivo* widefield imaging paradigm to visualize spontaneous neural activity in the superior colliculus (SC) of unanesthetized mouse pups.

(B) Calcium transients in the SC, color coded by time of occurrence, in a P7 control (*Tmem16a^fl/fl^;Snap25-T2A-GCaMP6s*) and P7 *Tmem16a* cKO (*Tecta-Cre;Tmem16a^fl/fl^;Snap25-T2A-GCaMP6s*) mouse.

(C) Spontaneous activation of SC regions of interest (ROIs) over a 10 minute acquisition in a P7 control and P7 *Tmem16a* cKO mouse.

(D) Quantification of SC wave frequency and wave duration. n = 7 control mice, 7 *Tmem16a* cKO mice; p = 0.3386, 0.3386 (frequency, duration), Wilcoxon rank sum test with Benjamini-Hochberg correction.

**Supplementary Figure 6. Sporadic but extensive activation of neurons and neuropil in the inferior colliculus with loss of cochlear TMEM16A, related to Figure 2**

(A) Schematic of two-photon imaging of spontaneous neural calcium transients within the inferior colliculus (IC) in pre-hearing mice.

(B) Representative spontaneous calcium transients within the left IC at a Z-depth of 150 μm in a P7 control (top,*Tmem16a^fl/fl^;Snap25-T2A-GCaMP6s*) and *Tmem16a* cKO mouse (bottom, *Tecta-Cre;Tmem16a^fl/fl^;Snap25-T2A-GCaMP6s*). Active grid region of interest (ROIs) are highlighted for a given spontaneous calcium transient. Astrocytes are labeled by topical application of SR101 (magenta) for image registration.

(C) Raster plot of calcium transients within 50 randomly selected ROIs within the field of view. Single events from left are identified by shading.

(D) Quantification of coordinated spontaneous event frequency in the IC. n = 4 colliculi from 3 control mice, 5 colliculi from 4 *Tmem16a* cKO mice; p = 0.0159, Wilcoxon rank sum test.

(E) Cumulative distribution of coordinated spontaneous event area. n = 275 events from 4 control colliculi (3 mice), 78 events from 5 *Tmem16a* cKO colliculi (4 mice); p = 2.6401e-4, two-sample Kolmogorov-Smirnov test.

**Supplementary Figure 7. Deletion of TMEM16A suppresses calcium transients in central auditory neurons throughout postnatal development, related to Figure 2**

(A) (left) Calcium transients in the IC from P10 control (*Tmem16a^fl/fl^;Snap25-T2A-GCaMP6s*) mouse. (right) Fluorescence trace over time of spontaneous activity in left (orange) and right (blue) IC with example single events highlighted.

(B) Same as (A), but from P10 *Tmem16a* cKO (*Tecta-Cre;Tmem16a^fl/fl^;Snap25-T2A-GCaMP6s*) mouse.

(C) Quantification of spontaneous event frequency. n = 6 control mice, 5 *Tmem16a* cKO mice; p = 0.0005, two-sample t-test with Benjamini-Hochberg correction for multiple comparisons.

(D) (left) Histogram of spontaneous event amplitude. (right) Cumulative distribution of spontaneous event amplitude. n = 1506 events from 6 control mice, 586 events from 5 *Tmem16a* cKO mice; p = 3.4338e-4, two-sample Kolmogorov-Smirnov test.

(E) Cumulative distribution of normalized band width (75^th^ percentile) for spontaneous events.

Inset indicates mean band width by animal. n = 674 events from 4 control mice, 235 events from 4 *Tmem16a* cKO mice; p = 3.3537e-27, two-sample Kolmogorov-Smirnov test. Inset: n = 4 control, 4 *Tmem16a* cKO mice; p = 0.0018, two-sample t-test.

**Supplementary Figure 8. Cochlear structure is preserved in *Tmem16a* cKO mice at maturity, related to Figure 3**

(A) Expression of TMEM16A in the cochlea at multiple developmental timepoints, as indicated by GFP (green) signal in *Tmem16a-GFP* mice. Hair cells are labeled by immunoreactivity to MyoVIIa (magenta).

(B) Hematoxylin and eosin stain of cross sections of P21 control (*Tmem16a^fl/fl^*) and P21 *Tmem16a* cKO (*Tecta-Cre;Tmem16a^fl/fl^*) basal cochleae. Black square indicates region of high magnification.

(C) Hair cells in whole mount preparations of apical, middle, and basal P21 control and *Tmem16a* cKO cochleae labeled by immunoreactivity to MyoVIIa (magenta).

(D) Quantification of inner hair cell (IHC) and outer hair cell (OHC) density. n = 4 cochleae from 4 control mice, 5 cochleae from 5 *Tmem16a* cKO mice; p = 0.0838, 0.0510 (IHCs, OHCs), linear mixed model.

(E) High magnification images of spiral ganglion neuron (SGN) soma, labeled by immunoreactivity to β-III tubulin (Tuj1, cyan), in apical, middle, and basal regions of the cochlea in P21 control and *Tmem16a* cKO mice.

(F) Quantification of SGN density. n = 4 cochleae from 4 control mice, 4 cochleae from 4 *Tmem16a* cKO mice; p = 0.4363, linear mixed model.

(G) Mean auditory brainstem response (ABR) trace to click (left) or 24 kHz pure tone pip (right) stimuli from 90 to 20 dB sound pressure level (SPL) in P21 control and *Tmem16a* cKO mice.

(H) Quantification of ABR threshold to click and pure tone stimuli. n = 11 control mice, 8 *Tmem16a* cKO mice; p = 0.3369, linear mixed model.

(I) Quantification of click ABR wave 1 (P1:N1) amplitude across a range of sound levels. n = 11 control mice, 8 *Tmem16a* cKO mice; mean ± SEM, p = 0.007877, linear mixed model.

**Supplementary Figure 9. Increased IC pure tone spatial activation across sound intensity levels following spontaneous activity disruption, related to Figure 4.**

(A) Tone-evoked neural calcium transients in IC from P14 control (*Tmem16a^fl/fl^;Snap25-T2A-GCaMP6s*) and P14 *Tmem16a* cKO (*Tecta-Cre;Tmem16a^fl/fl^;Snap25-T2A-GCaMP6s*) mice at 80 dB SPL.

(B) Enlarged merged image from white square in (A) shows tonotopic segregation of pseudocolored pure tone responses; cyan and yellow colors indicate areas of overlap.

(C) Measurement of 6 and 12 kHz evoked band width (75^th^ percentile) normalized to peak fluorescence response amplitude along the tonotopic axis of the IC at 80 dB SPL in P14 control and *Tmem16a* cKO mice.

(D) Quantification of spatial evoked fluorescence by frequency along tonotopic axis at 80 dB SPL. n = 9 control mice, 8 *Tmem16a* cKO mice; mean ± SEM, p = 0.0078, linear mixed model with Sidák post hoc test.

(E) Quantification of mean band width by mouse across all tested frequencies at 80 dB SPL. n = 9 control mice, 8 *Tmem16a* cKO mice; p = 0.0045, two-sample t-test.

(F) Same as (A), but at 60 dB SPL.

(G) Same as (B), but at 60 dB SPL.

(H) Measurement of 6 and 12 kHz pure tone evoked band width (75^th^ percentile) normalized to peak fluorescence response amplitude along the tonotopic axis of the IC at 60 dB SPL in control and *Tmem16a* cKO mice.

(I) Quantification of spatial evoked fluorescence along tonotopic axis at 60 dB SPL. n = 8 control mice, 7 *Tmem16a* cKO mice; mean ± SEM, p = 0.0053, linear mixed model with Sidák post hoc test.

(J) Quantification of mean band width by mouse across all tested frequencies at 60 dB SPL. n = 8 control mice, 7 *Tmem16a* cKO mice; p = 1.6523e-4, two-sample t-test.

**Supplementary Figure 10. Enhanced sound-evoked calcium responses of colliculus neurons with disruption of pre-hearing spontaneous activity, related to Figure 5**

(A) Mean maximum fluorescence changes within responsive neuronal soma elicited by increasing intensity of a 3 kHz pure tone within P14 control (*Tmem16a^fl/fl^;Snap25-T2A-GCaMP6s*) and P14 *Tmem16a* cKO (*Tecta-Cre;Tmem16a^fl/fl^;Snap25-T2A-GCaMP6s*) inferior colliculus (IC). n = 55 cells from 4 control mice, 88 cells from 5 *Tmem16a* cKO mice; mean ± SEM, p = 0.0347, linear mixed model with Sidák post hoc test.

(B) Same as (A), but for 6 kHz pure tone. n = 59 cells from 6 control mice, 79 cells from 8 *Tmem16a* cKO mice; p = 0.0042, linear mixed model with Sidák post hoc test.

(C) Same as (A), but for 12 kHz pure tone. n = 14 cells from 3 control mice, 27 cells from 3 *Tmem16a* cKO mice; p = 0.0116, linear mixed model with Sidák post hoc test.

(D) Maximum amplitude of tone-evoked neural soma fluorescence changes to 3 kHz, 6 kHz, 9.5 kHz, and 12 kHz stimuli at any sound level. n = 55, 59, 30, 14 control cells (3 kHz, 6 kHz, 9.5 kHz, 12 kHz), 88, 79, 77, 27 *Tmem16a* cKO cells; p = 1.2909e-7, 1.7186e-4, 4.7945e-4, 0.0062 (3 kHz, 6 kHz, 9.5 kHz, 12 kHz), Wilcoxon rank sum test.

(E) (top) Assessment of 9.5 kHz evoked fluorescence responses in neuropil by exclusion of all neural soma (black ROIs masking image). (bottom) Mean fluorescence changes in neuropil averaged across the field of view at 3 different sound pressure levels. n = 5 control mice, 6 *Tmem16a* cKO mice, mean ± SEM.

(F) Mean maximum fluorescence changes of neuropil in response to increasing 9.5 kHz pure tone stimulus intensity. n = 5 control mice, 6 *Tmem16a* cKO mice; mean ± SEM, p = 1.5970e-4, linear mixed model.

(G) Plot of maximum evoked calcium amplitudes of all tested frequencies at 70 dB SPL overlaid with a single term Gaussian fit to calculate bandwidth (full width half max of Gaussian fit, indicated by black horizontal line).

(H) Cumulative distribution of maximum Gaussian bandwidth at 70 dB SPL. n = 159 cells (8 mice), 227 *Tmem16a* cKO cells (7 mice), p = 1.1961e-20, two-sample Kolmogorov-Smirnov test.

(I) (left) Pseudocolored tone-evoked fluorescence in IC neurons at 70 dB SPL in a control and *Tmem16a* cKO mouse. White arrows indicate neurons responsive to both stimuli in merged image. (right) Quantification of maximum frequency bandwidth of IC neurons at 70 dB SPL. n = 207 control cells (8 mice), 252 *Tmem16a* cKO cells (7 mice); p = 1.1137e-54, two-sample Kolmogorov-Smirnov test.

(J) Mean fluorescence change in response to a neuron’s best frequency stimulus at 80 to 50 dB SPL. Vertical bar indicates tone presentation. n = 235 control cells, 295 *Tmem16a* cKO cells; mean ± SEM.

**Supplementary Figure 11. Spatial compression of frequency representation is present across a range of sound intensity levels, related to Figure 6**

(A) Individual and pseudocolored merged images of 80 dB sound pressure level (SPL) tone-evoked neural calcium transients in IC from a P14 control (*Tmem16a^fl/fl^;Snap25-T2A-GCaMP6s*) and P14 *Tmem16a* cKO (*Tecta-Cre; Tmem16a^fl/fl^;Snap25-T2A-GCaMP6s*) mouse. Rectangular ROIs were placed along the tonotopic axis of the IC (low to high frequency) to determine the location of best frequency (BF).

(B) (left) Plot of mean tone-evoked fluorescence along the tonotopic axis of the IC at 80 dB SPL. Dashed line indicates location of peak response, grey shading depicts shift in peak response location between control and cKO mice. (right) Same as (left) but normalized to maximum amplitude. n = 9 control mice, 8 *Tmem16a* cKO mice; mean ± SEM.

(C) Quantification of peak response location to 80 dB SPL stimuli relative to 3 kHz (lowest frequency) peak response location. n = 9 control mice, 6 *Tmem16a* cKO mice; mean ± SEM, p = 0.00169, linear mixed model.

(D) Same as (A), but at 60 dB SPL. Increased gain of merged image enables visualization of distinct frequency domains.

(E) (left) Plot of mean tone-evoked fluorescence along the tonotopic axis of the IC at 60 dB SPL. Control and *Tmem16a* cKO profiles were aligned to 6 kHz responses (lowest suprathreshold frequency with consistent responses across animals at 60 dB SPL). (right) Same as (left) but normalized to maximum amplitude. n = 5 control mice, 5 *Tmem16a* cKO mice, mean ± SEM.

(F) Quantification of peak response location of increasing frequency stimuli at 60 dB SPL relative to 6 kHz peak response location. n = 5 control mice, 5 *Tmem16a* cKO mice; mean ± SEM, p = 0.00263, linear mixed model.

(G) Plot of mean normalized tone-evoked fluorescence along the tonotopic axis of the IC to 6 kHz and 12 kHz pure tones at 80 and 60 dB SPL. Dashed line indicates peak response location, shading indicates difference between 6 kHz and 12 kHz peak location. n = 5-9 control mice, 5-8 *Tmem16a* cKO mice.

## Supplementary Movie captions

**Supplementary Movie 1. TMEM16A channels are required for coordinated IHC activation, related to Supplementary** Figure 3

Confocal imaging of calcium transients in inner hair cells (IHCs) and inner supporting cells (ISCs) in P7 control (*Tecta-Cre;Tmem16a^fl/+^;R26-lsl-GCaMP3*, top) and *Tmem16a* cKO (*Tecta-Cre;Tmem16a^fl/fl^;R26-lsl-GCaMP3*, bottom) excised cochlea expressing the genetically encoded calcium indicator GCaMP3. Images were collected at 2 Hz, playback is 10 frames per second.

Supplementary Movie 2. Deletion of TMEM16A suppresses calcium transients within the inferior colliculus, related to Figure 2

Wide-field imaging of spontaneous neural activity in the inferior colliculus of P7 control (*Tmem16a^fl/fl^;Snap25-T2A-GCaMP6s*, top) and *Tmem16a* cKO (*Tecta-Cre;Tmem16a^fl/fl^;Snap25-T2A-GCaMP6s*, bottom) mice that expressed GCaMP6s pan-neuronally under the *Snap25* promoter. Images were collected at 10 Hz, playback is 20 frames per second.

Supplementary Movie 3. Increased gain of pure tone neuronal responses following disruption of pre-hearing spontaneous activity, related to Figure 5

Two photon imaging of sound-evoked neural activity in the inferior colliculus of P14 control (*Tmem16a^fl/fl^;Snap25-T2A-GCaMP6s*, left) and *Tmem16a* cKO (*Tecta-Cre;Tmem16a^fl/fl^;Snap25-T2A-GCaMP6s*, right) mice. 6 kHz amplitude modulated tones are presented at a peak of 93, 73, and 53 dB SPL for 200 ms to the contralateral ear. Images were collected at 5 Hz, playback is 3 frames per second.

Supplementary Movie 4. Broader frequency receptive fields in IC neurons with spontaneous activity disruption, related to Figure 5

Two photon imaging of sound-evoked neural activity in the inferior colliculus of P14 control (*Tmem16a^fl/fl^;Snap25-T2A-GCaMP6s*, left) and *Tmem16a* cKO (*Tecta-Cre;Tmem16a^fl/fl^;Snap25-T2A-GCaMP6s*, right) mice. 4.4 kHz (green) and 7.8 kHz (magenta) amplitude modulated tones are presented at 70 dB SPL for 200 ms to the contralateral ear. White indicates cells and processes responsive to both frequencies, with increased overlap of pure tone responsive domains in *Tmem16a* cKO mice. Images were collected at 5 Hz, playback is 2 frames per second.

Supplementary Movie 5. Spatial compression of frequency representation with spontaneous activity disruption, related to Figure 6

Widefield imaging of pseudocolored sound-evoked neural activity in the right inferior colliculus of P14 control (*Tmem16a^fl/fl^;Snap25-T2A-GCaMP6s*, left) and *Tmem16a* cKO (*Tecta-Cre;Tmem16a^fl/fl^;Snap25-T2A-GCaMP6s*, right) mice. 3 kHz (green) and 9.5 kHz (magenta) amplitude modulated tones are presented at a peak of 100 dB SPL for 1 s to the contralateral ear. Images were collected at 10 Hz, playback is 3 frames per second.

