## Supplementary Figures for "Developmental spontaneous activity promotes sensory domains, frequency tuning and proper gain in central auditory circuits"

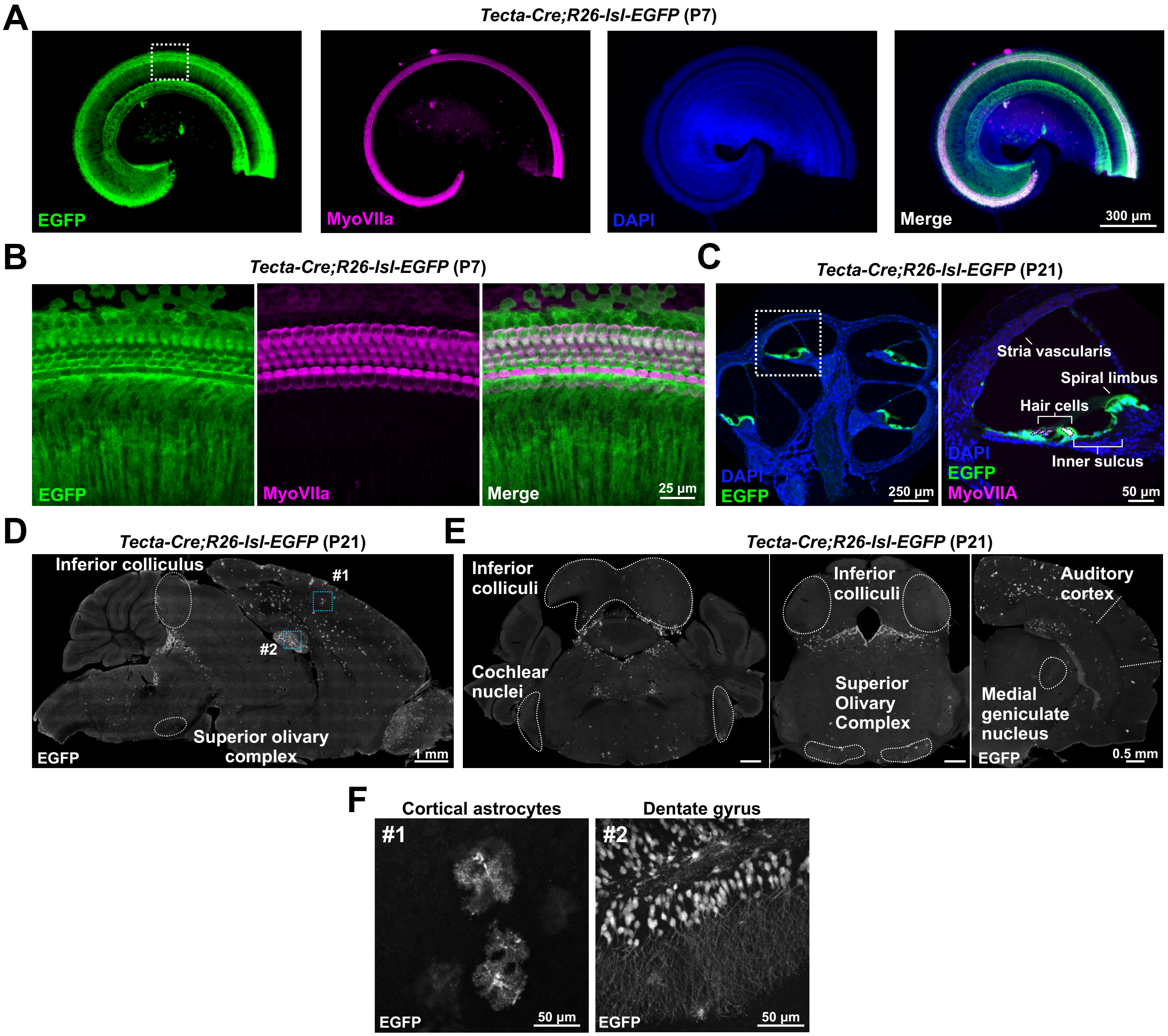


### Supplementary Figure 1

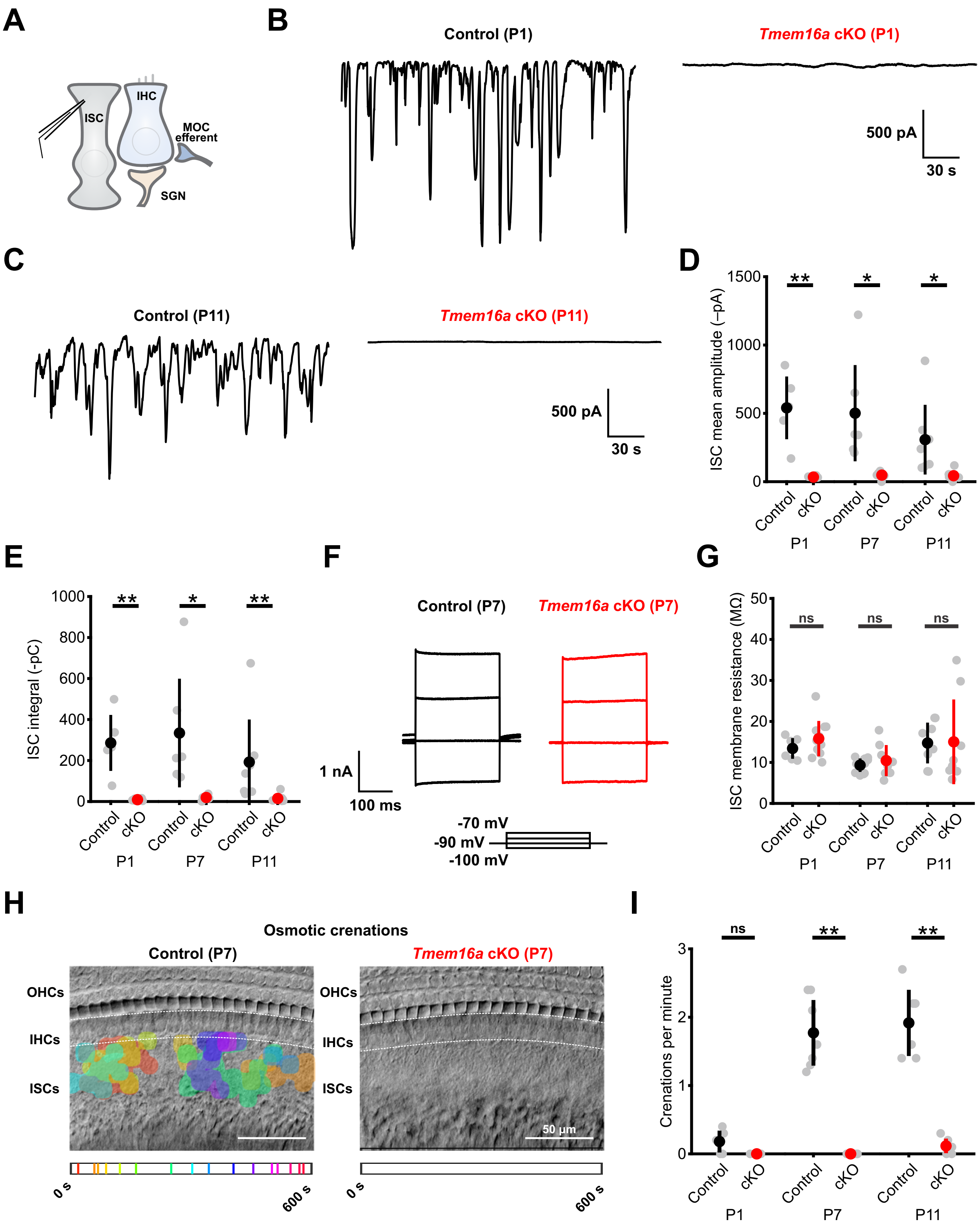


### Supplementary Figure 2

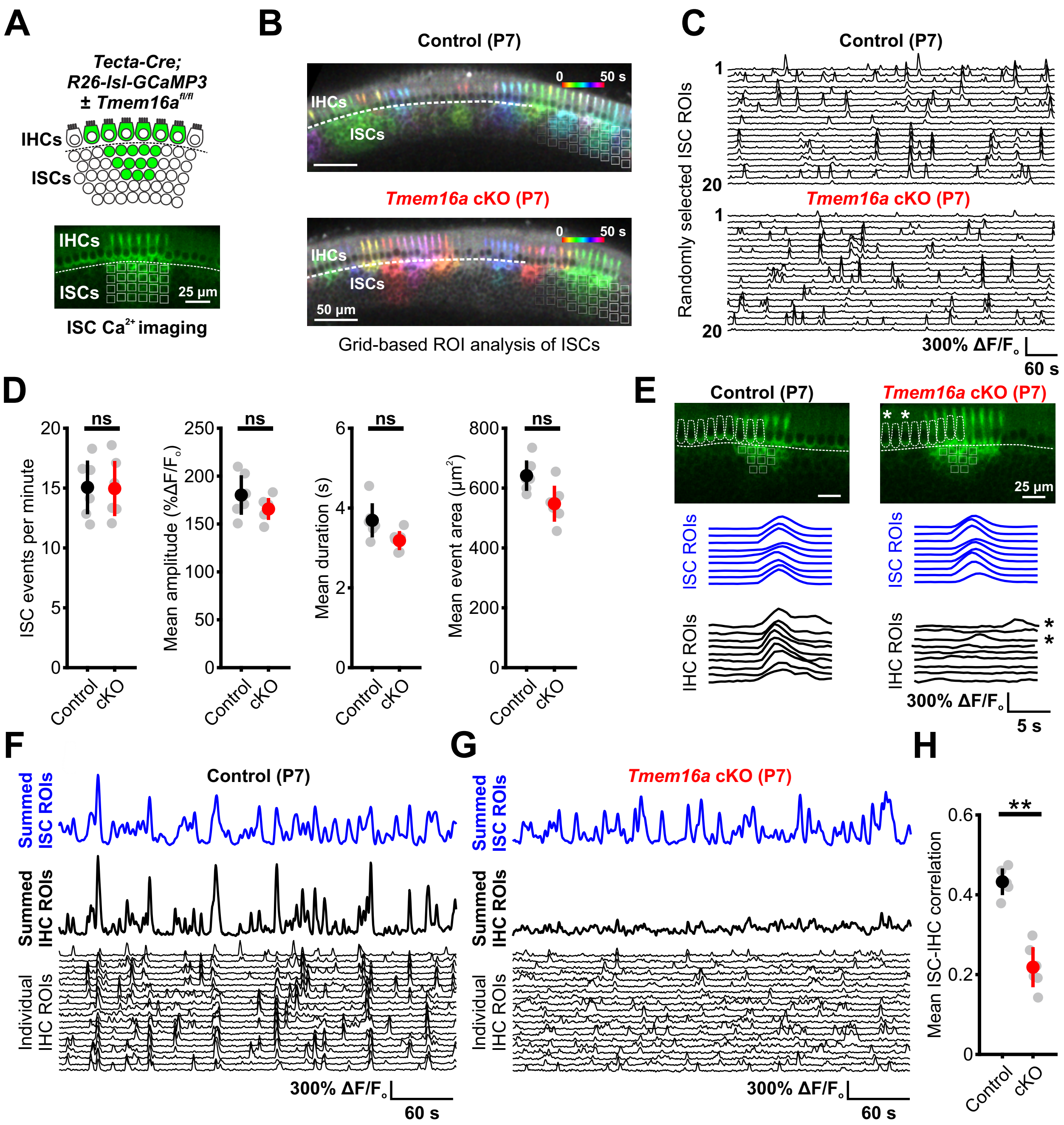


### Supplementary Figure 3

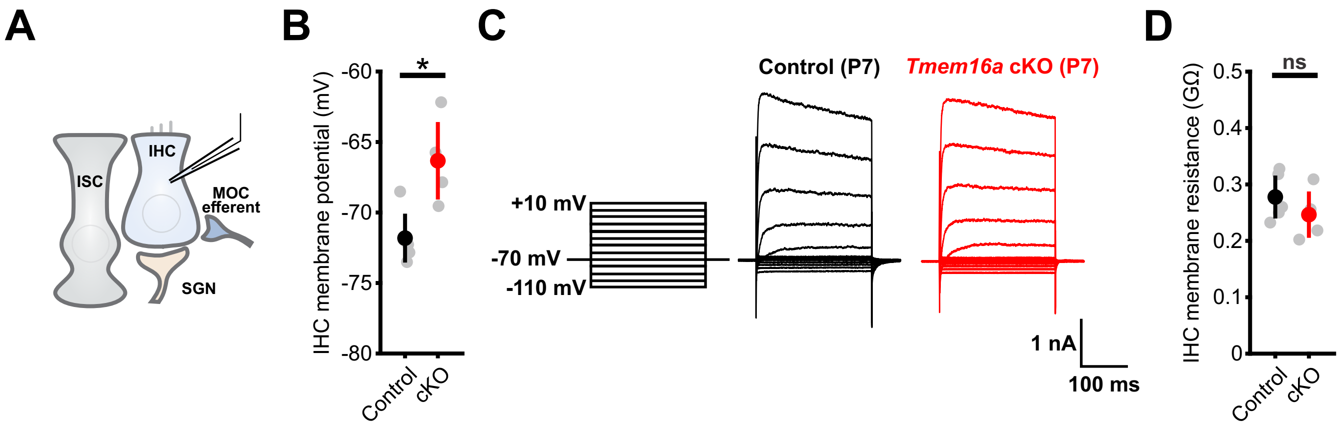


**Supplementary Figure 4**

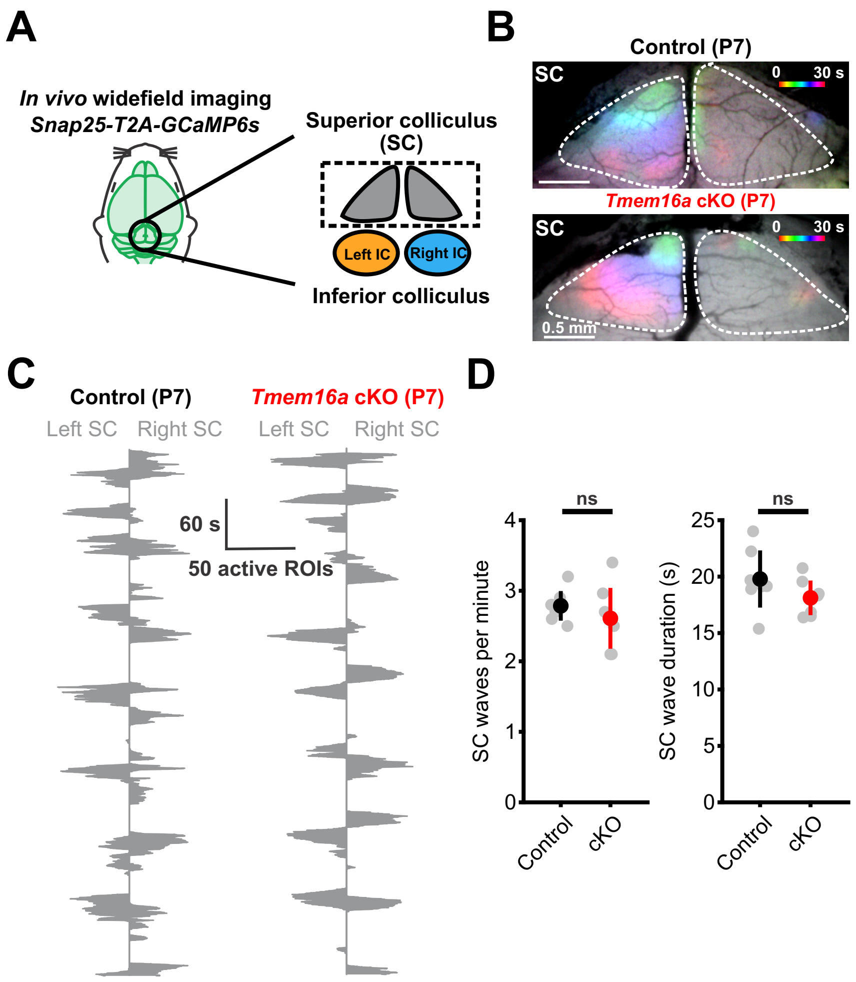


### Supplementary Figure 5

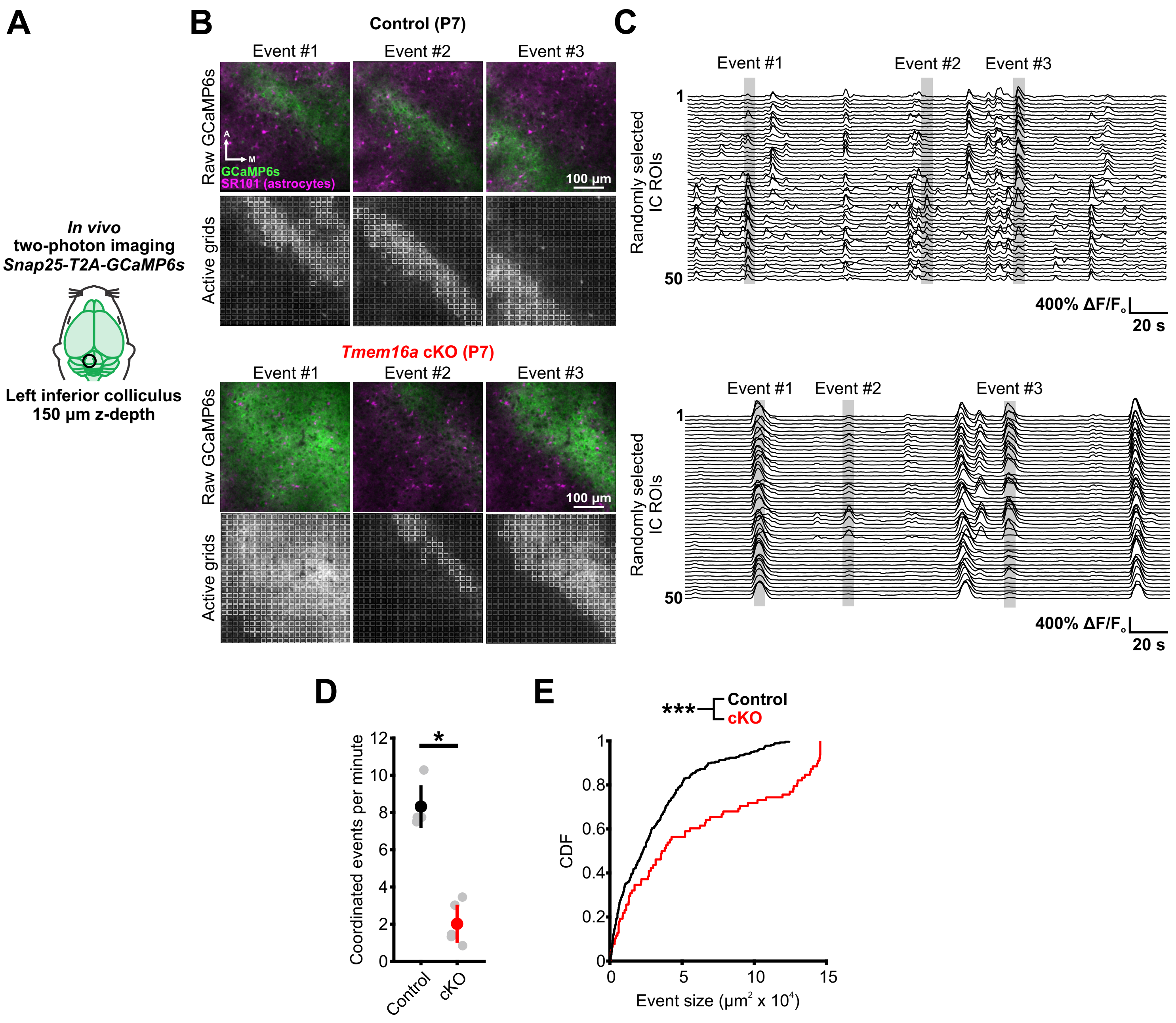


### Supplementary Figure 6

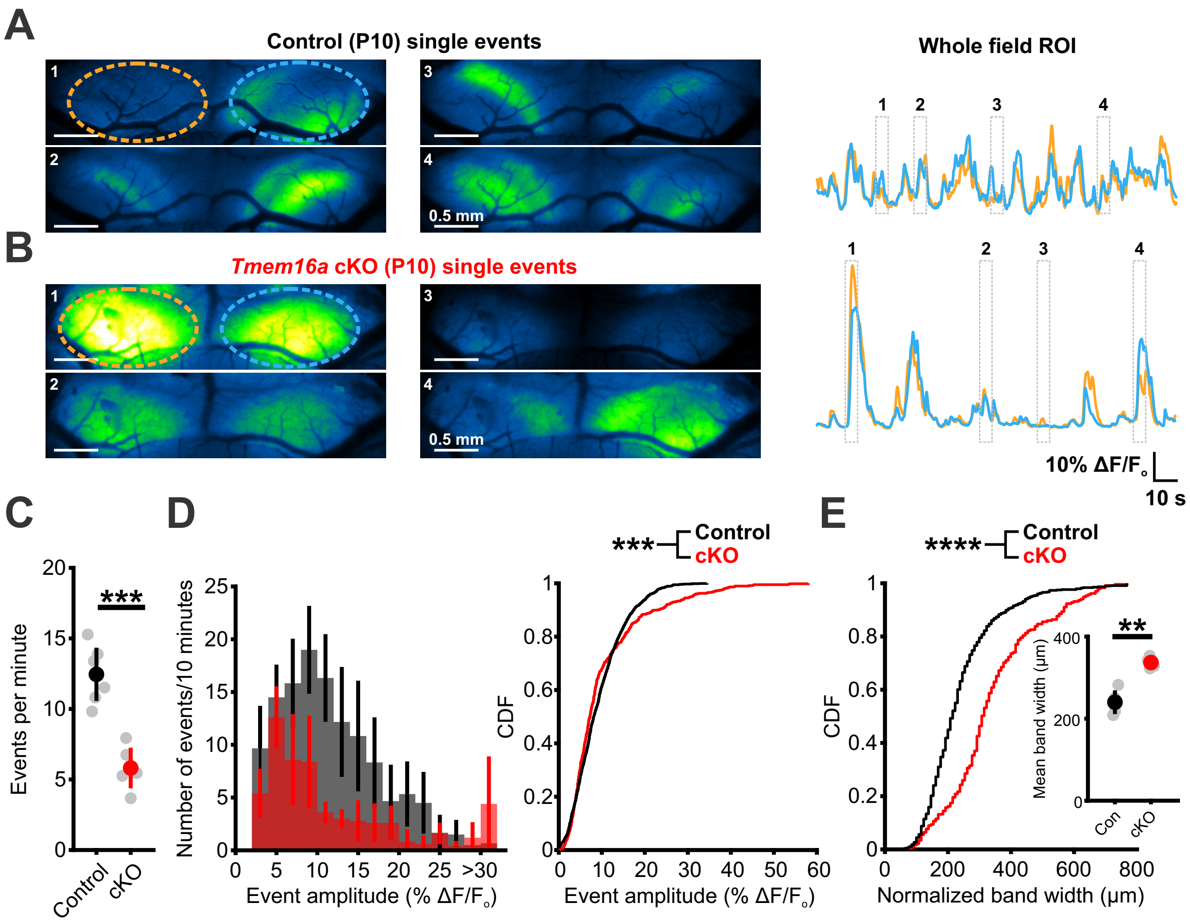


### Supplementary Figure 7

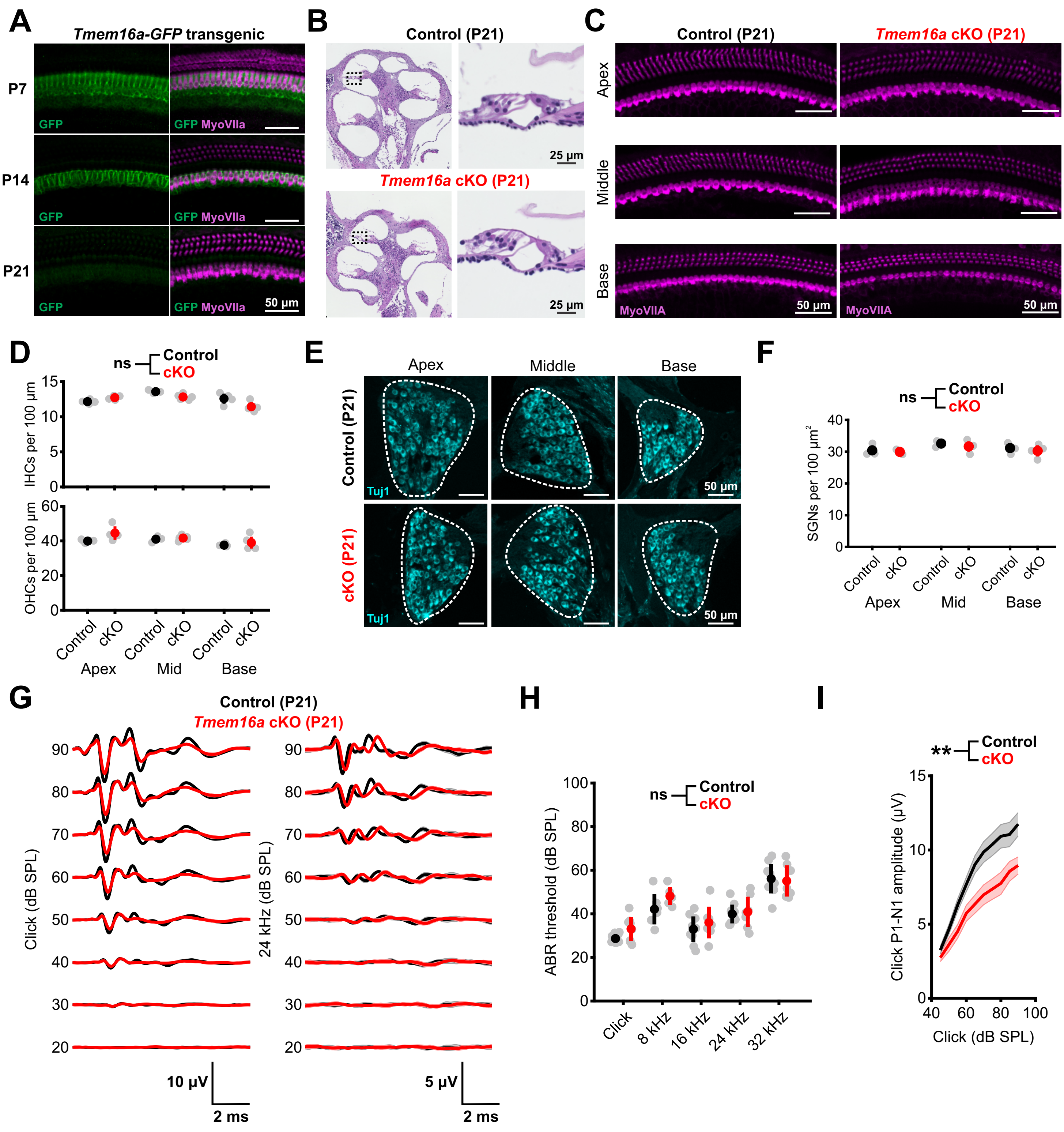


### Supplementary Figure 8

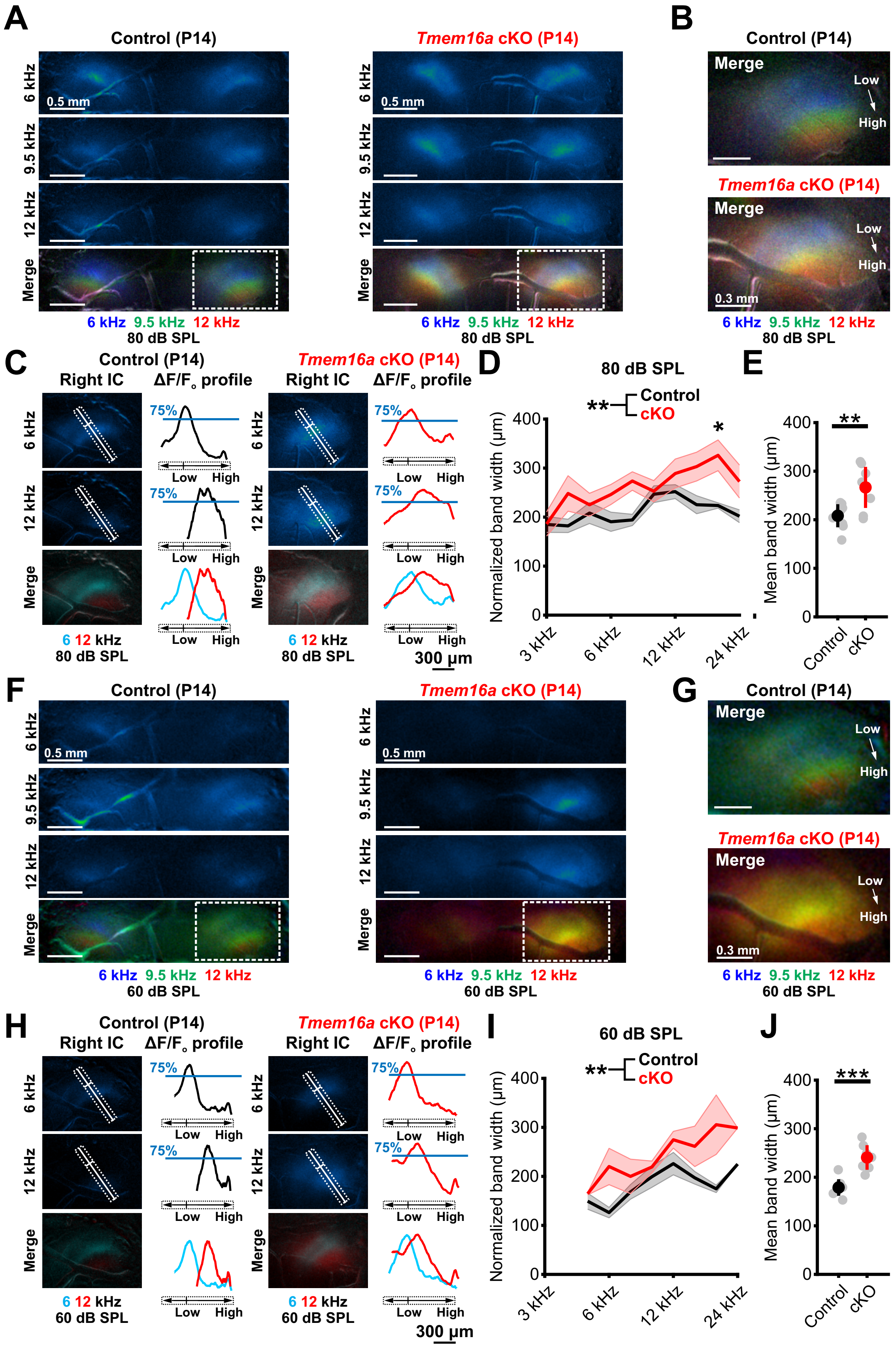


### Supplementary Figure 9

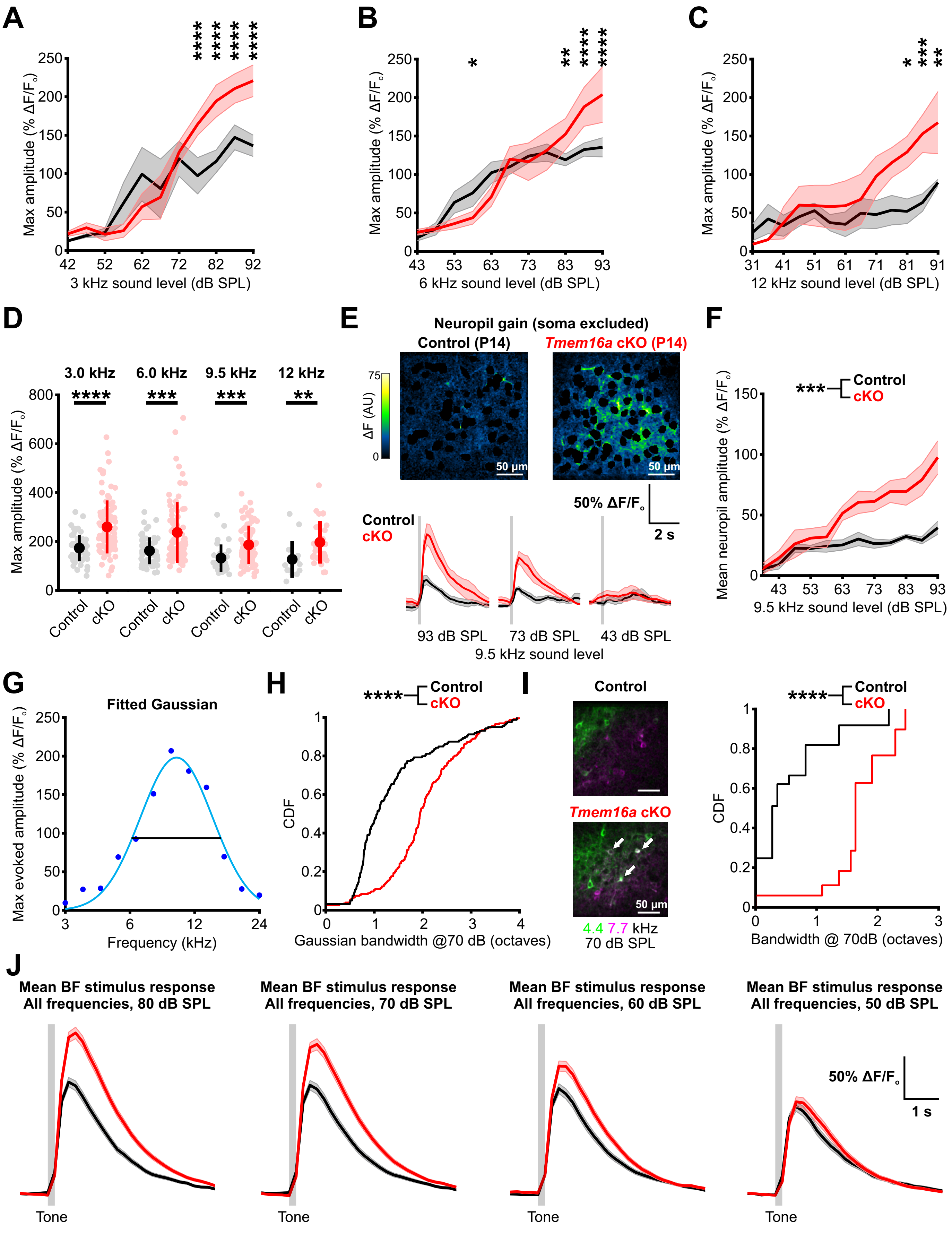


### Supplementary Figure 10

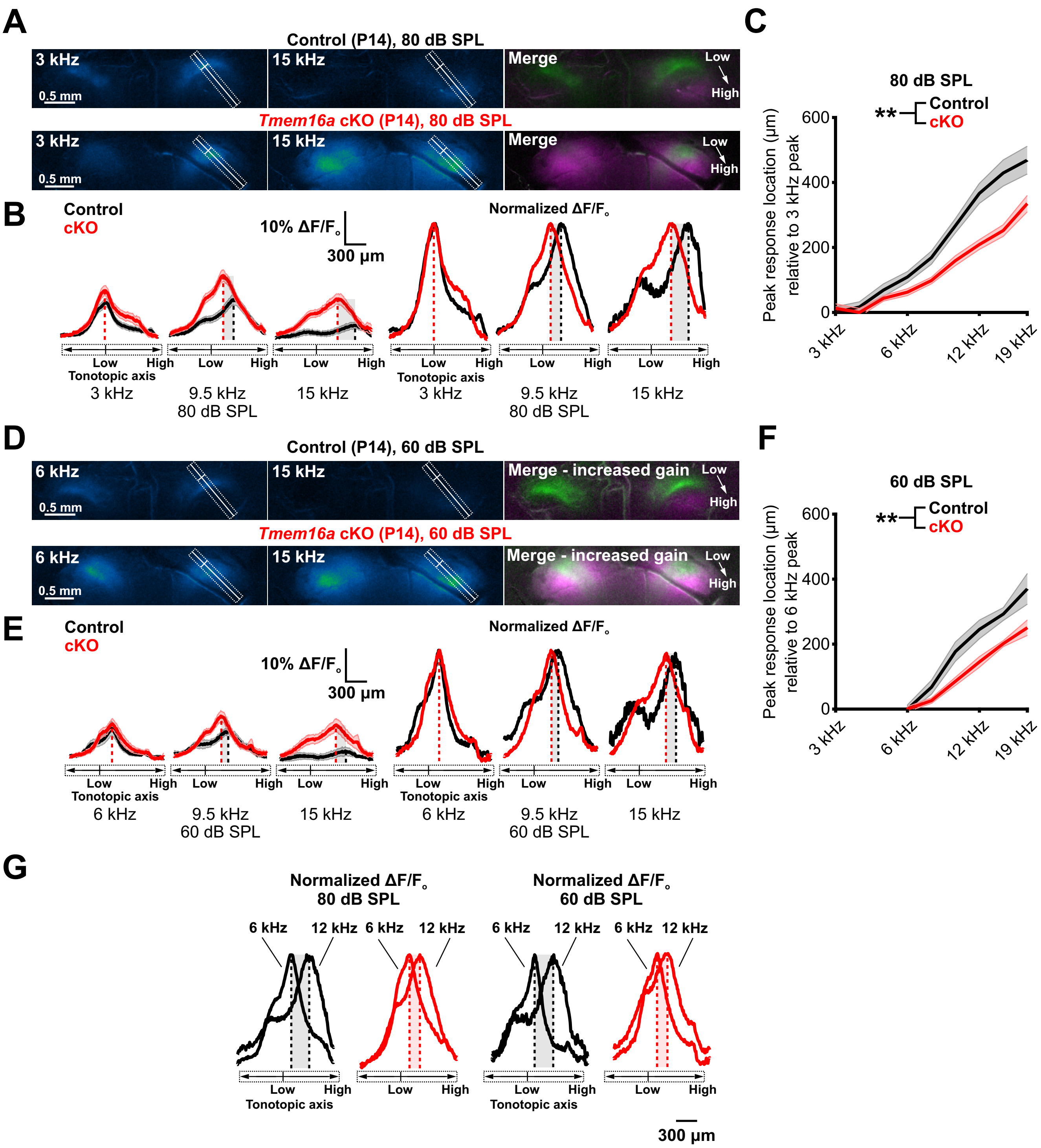


### Supplementary Figure 11
